# A secondary metabolite drives intraspecies antagonism in a gut symbiont that is inhibited by cell wall acetylation

**DOI:** 10.1101/2021.06.11.448121

**Authors:** Mustafa Özçam, Jee-Hwan Oh, Restituto Tocmo, Deepa Acharya, Shenwei Zhang, Theresa Astmann, Mark Heggen, Silvette Ruiz-Ramírez, Fuyong Li, Christopher C. Cheng, Eugenio Vivas, Federico E. Rey, Jan Claesen, Tim Bugni, Jens Walter, Jan-Peter van Pijkeren

## Abstract

The mammalian microbiome encodes numerous secondary metabolite biosynthetic gene clusters, yet their role in microbe-microbe interactions is unclear. Here, we characterized two polyketide synthase gene clusters (*fun* and *pks*) in the gut symbiont *Limosilactobacillus reuteri.* The *pks*, but not the *fun* cluster, encodes antimicrobial activity. Forty-one out of 51 *L. reuteri* strains tested are sensitive to Pks products, which was independent of strains’ host origin. The sensitivity to Pks was also established in intraspecies competition experiments in gnotobiotic mice. Comparative genome analyses between Pks-resistant and sensitive strains identified an acyltransferase gene (*act*) that is unique to Pks-resistant strains. Subsequent cell wall analysis of the wild-type and the *act* mutant strains showed that Act acetylates cell wall components. The *pks* mutants lost their competitive advantage and *act* mutants lost their Pks resistance *in vivo*. Thus, our findings provide insight into how closely related gut symbionts can compete and co- exist in the gastrointestinal tract.

## INTRODUCTION

The mammalian gastrointestinal tract is inhabited by trillions of microorganisms that coexist with their host (Sender et al., 2016). While factors like host genetics, immune status, nutritional resources and colonization history affect microbial composition (Bonder et al., 2016; David et al., 2014; Hooper et al., 2015; Martínez et al., 2018; Snijders et al., 2016; Turnbaugh et al., 2009; Zarrinpar et al., 2014), the relationship between microbes is determined by competitive interactions (Boon et al., 2014). Bacteria have developed numerous strategies that mediate survival and competition, including the production of broad and narrow spectrum antimicrobials (Sassone-Corsi et al., 2016). On the other hand, acquisition of resistance genes and/or modification of the cell wall can help bacteria survive the antimicrobial warfare in the gut (Murray and Shaw, 1997).

Some bacterial secondary metabolites have antimicrobial activity against other community members. Their interactions—known as interference competition—are important in the assembly and the maintenance of microbial communities (Jacobson et al., 2018). Most secondary metabolites are produced by biosynthetic gene clusters (BGCs) and polyketide synthase (PKS) gene clusters form a prominent subclass, involved in the biosynthesis of carbon chain backbones from the repeated condensation of acyl-CoA building blocks (Lin et al., 2015; Medema et al., 2014). A survey of 2,430 reference genomes from the Human Microbiome Project uncovered more than 3,000 BGCs (Donia et al., 2014). So far, only seven PKS-like BGCs encoded by gut-associated bacteria have been functionally characterized (Figure S1).

As a class of polyketide synthase products, aryl polyenes are lipids with an aryl head group conjugated to a polyene tail (Lin et al., 2015) and are widely distributed in soil and host-associated bacteria (Cimermancic et al., 2014; Youngblut et al., 2020). Although, many polyene compounds isolated from terrestrial and marine microbes possess antimicrobial effects *in vitro* (Herbrík et al., 2020; Lee et al., 2020; Li et al., 2021; Zhao et al., 2021), virtually nothing is known about their ecological role in microbe-host and microbe-microbe interactions in the mammalian gastrointestinal tract (Aleti et al., 2019). While metagenome studies are critically important to identify novel polyene-like BGCs within the microbiome (Hiergeist et al., 2015; Medema et al., 2011), the assessment of model organisms and their isogenic mutants in the appropriate ecological context is critical to advance our knowledge on the biological function of these BGC-derived compounds.

*Limosilactobacillus reuteri*, until recently known as *Lactobacillus reuteri* (Zheng et al., 2020), is a gut symbiont species that inhabits the gastrointestinal tract of various vertebrates, including rodents, birds and primates (Duar et al., 2017). This, combined with the available genome editing tools (Oh and Van Pijkeren, 2014; Van Pijkeren et al., 2012; Zhang et al., 2018), make *L. reuteri* an ideal model organism to study microbe-microbe, microbe-phage and microbe-host interactions (Lin et al., 2018; Oh et al., 2019; Özçam et al., 2019; Walter et al., 2011).

Previously, we screened a *L. reuteri* strain library derived from different hosts and identified that *L. reuteri* R2lc activates the aryl hydrocarbon receptor (AhR), a ligand activated transcription factor that plays an important role in mucosal immunity by inducing the production of interleukin-22 (Zelante et al., 2013). By whole genome sequencing, comparative genomics, and targeted gene deletion, we identified that R2lc harbors two genetically distinct PKS clusters (*pks* and *fun*), each encoded from a multi-copy plasmid. We demonstrated that the *pks* but not the *fun* cluster is responsible for R2lc mediated AhR activation (Özçam et al., 2019)

In this study, we found that the *pks* cluster in *L. reuteri* R2lc also encodes antimicrobial activity. Intraspecies competition experiments in gnotobiotic mice with the wild-type and the *pks* deletion mutant revealed that Pks expression can provide a competitive advantage. Remarkably, a small number *L. reuteri* strains were resistant to the antimicrobial activity of Pks. We discovered this resistance was driven by a gene encoding an acyltransferase that increases acetylation of the cell wall Thus, our findings uncovered mechanisms by which gut symbionts can compete and co-exist with closely related strains through secondary metabolite production and cell-wall modification.

## RESULTS

### *L. reuteri* Pks inhibits the competitor strain

The rodent isolate *L. reuteri* R2lc contains two PKS clusters, *fun* and *pks* (Figure 1A). In both the human and rodent gut ecosystem, *L. reuteri* R2lc outcompetes most *L. reuteri* strains, including the rodent isolate *L. reuteri* 100-23 (Duar et al., 2017; Oh et al., 2009). Because select PKS products have antimicrobial activity (Lin et al., 2015), we tested to what extent Pks and Fun provide strain R2lc with a competitive advantage. We performed *in vitro* competition experiments in batch cultures using 1:1 mixtures of the rodent gut isolates *L. reuteri* 100-23 and

**Figure 1.**
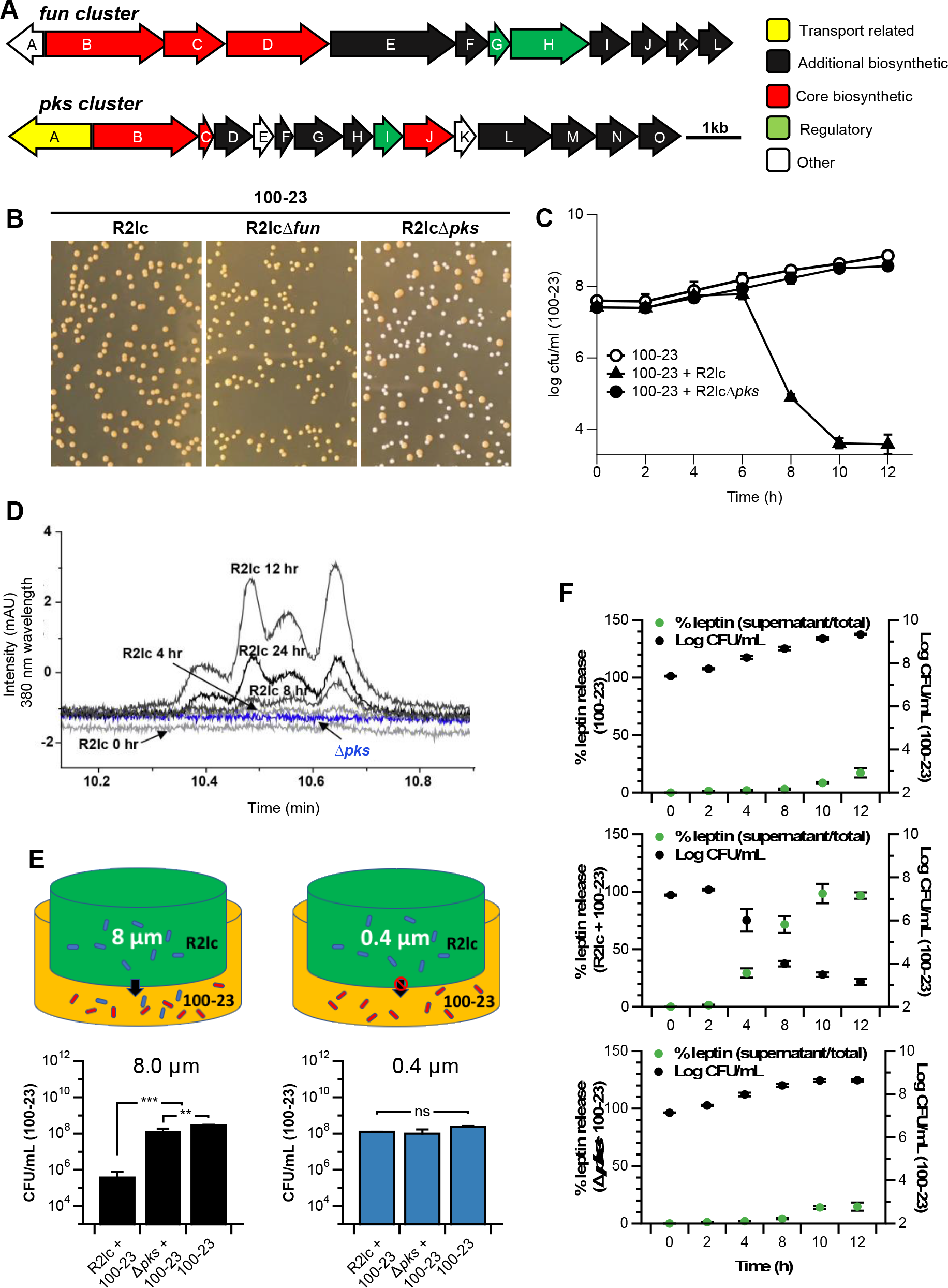
A polyketide synthase cluster in *L. reuteri* R2lc produces polyene-like compounds that lyse competitor strains. **A)** The *fun* cluster (top) spans 13.4 kb containing 12 Open Reading Frames (ORFs), and the *pks* cluster (bottom) spans 11.3 kb containing 15 ORFs. Transport-related, additional biosynthetic, regulatory, and other genes are represented by different colors. **B)** R2lc and R2lcΔ*fun* but not R2lcΔ*pks* inhibits *L. reuteri* 100-23. **C)** R2lc has bactericidal effect against 100-23. Single culture (OD_600_=0.1) or co-cultures (OD_600_=0.05 from each strain) were mixed and incubated in MRS broth (pH: 4.0, 37°C) and samples were collected every two hours for up to 12 hours. See also Figure S2. **D)** UPLC-PDA-MS analysis of R2lc and Δ*pks* mutant. R2lc but not Δ*pks* produces unique compounds with a maximum absorption of 380nm (Black: R2lc, blue: Δ*pks*). See also Figure S3. **E)** R2lc-Pks-mediated killing requires cell-cell interaction. Growth (24h) of R2lc and 100-23 separated by a 8 µm filter resulted in killing of 100-23 in a Pks-dependent manner (top left schematic; black bars) while separation by a 0.4 µm filter (top right schematic; blue bars) did not affect 100-23 survival, as determined by viable plate count. ns. no statistical significance (p > 0.05); **p < 0.01; ***p < 0.001. See also Figure S4. **F)** R2lc-Pks lyses competitor strain 100-23. Strain 100-23 was engineered to accumulate recombinant leptin intracellularly. Percentage leptin released (primary y-axis, green) and bacterial counts of 100-23 (secondary y-axis, black) were determined for a monoculture of 100-23 (top panel) and co-cultures of R2lc+100- 23 (middle panel) and R2lcΔ*pks*+100-23 (bottom panel). All data shown are means ± SD and are based on three biological replicates.

*L. reuteri* R2lc wild type, or our previously generated PKS mutants (R2lcΔ*fun* or R2lcΔ*pks*) (Özçam et al., 2019). Following 24 hours of incubation, the mixed cultures were plated. On agar, R2lc and its derivatives form pigmented colonies, while 100-23 colonies are opaque in color. Co- incubation of R2lc + 100-23 or R2lcΔ*fun* + 100-23 only recovered pigmented colonies; however, co-incubation of R2lcΔ*pks* + 100-23 yielded a mixture of pigmented and opaque-colored colonies (% pigmented:opaque colony distribution is 29:71) (Figure 1B). These data suggest that the *pks* gene cluster, but not the *fun* gene cluster, provides strain R2lc with a competitive advantage when co-cultured with *L. reuteri* 100-23.

### *L. reuteri* Pks has a time-dependent bactericidal effect

To understand the dynamics by which R2lc outcompetes 100-23, we performed a time course competition experiment in batch cultures. Mixtures (1:1, OD600 = 0.05 per strain) of R2lc + 100-23, and R2lcΔ*pks* + 100-23 were prepared. As a control, a monoculture of 100-23 was grown. We harvested samples every two hours and determined the CFU levels by standard plate count. We observed that R2lc and R2lcΔ*pks* have a similar growth dynamic (μmax of R2lc= 0.19±0.02, μmax of R2lcΔ*pks*= 0.16±0.003, *p* = 0.18), and 100-23 grows slightly faster compared to R2lc and R2lcΔ*pks* in mono-culture (Figure S2A). Up to six hours, mixtures of R2lc + 100-23 and R2lcΔ*pks* + 100-23 yielded similar levels of 100-23 colonies, which were comparable to the levels obtained when 100-23 was cultured independently. However, after eight and ten hours of co-incubation of R2lc + 100-23, 100-23 CFU levels were reduced by three and five orders of magnitude, respectively, while 100-23 continued to grow in the R2lcΔ*pks* + 100-23 mixture (Figure 1C). Importantly, the sharp decline in 100-23 CFU counts suggests Pks elicits a strong bactericidal activity against 100-23. Also, the fact that killing of 100-23 is initiated after 6-hours of co-culture suggests that production of Pks may be growth phase dependent.

In mice, *L. reuteri*—including strain 100-23—form a biofilm on the stratified squamous epithelium of the forestomach (Frese et al., 2013; Lin et al., 2018; Savage et al., 1968). Cell numbers of 100-23, R2lc or R2lcΔ*pks* in gastric biofilms were similar (4.3×10^8^ ± 2.6×10^8^, 2.0×10^8^ ± 3.0×10^7^, 1.7×10^8^ ± 4.1×10^7^ CFU/ml, respectively) (Figure S2A). In biofilm co-cultures, however, R2lc was 750-fold more abundant than 100-23 (*p* = 0.002). The ability of R2lc to outcompete 100-23 in biofilm is mediated by Pks, because co-incubation of R2lcΔ*pks* with 100-23 recovered similar levels of CFUs for both strains (*p* = 0.27) (Figure S2B).

### *L. reuteri pks* cluster produces unique polyene-like compounds

To gain more insight into *L. reuteri* R2lc Pks production *in vitro*, we performed Ultra Performance Liquid Chromatography coupled with Photodiode Array and Mass Spectrometer (UPLC-PDA-MS) analysis. By comparing the chromatograms obtained from R2lc and R2lcΔ*pks* cultures, we found that R2lc produces a family of unique compounds with a maximum wavelength of 380 nm. These compounds eluted towards the end of the gradient run where the solvent composition was close to 100% methanol, indicating a hydrophobic characteristic. The large retention time of the compounds on the C18 column and the maximum absorption wavelength are consistent with previously identified polyene compounds (Gruber and Steglich, 2007). Therefore, the putative products of the *pks* cluster are predicted to be polyene compounds.

The UV chromatogram extracted from the R2lc culture at different time points shows an increase in the production of these compounds up to 12 hours of incubation, after which the intensity reduces as shown in samples at 24 hours (Figure 1D). We subsequently analyzed the Liquid Chromatography-Mass Spectrometry (LC-MS) data of R2lc and Δ*pks* samples collected above to identify the polyene products produced by R2lc. The mass spectrum of the peak between 10.52 min – 10.58 min in the R2lc chromatogram at 12 hours clearly showed an ion with *m/z* [M+H]^+^ value of 257.1172 and corresponding *m/z* [M+Na]^+^ value of 279.0092, which were absent in the 12-hour culture of *Δpks* (Figure S3). The high-resolution mass for the compound led to the predicted molecular formula of C16H16O3. The reduction of Pks compounds after 12 hours of incubation suggests instability of Pks molecules in our experimental setup.

Taken together, these data suggest that the antibacterial compound is produced in a time-dependent manner.

### Pks producing *L. reuteri* R2lc lyses competitor strain through cell-cell interaction

To further enhance our understanding of *L. reuteri* R2lc Pks-mediated killing, we performed an experiment where strains R2lc and 100-23 were cultured in individual chambers separated by a 0.4 µm or 8µm filter (Figure 1E, top schematic). The 0.4 and 8 µm membranes will allow flowthrough of small molecules, metabolites and secreted soluble proteins; only the 8 µm membrane allows flowthrough of bacteria. When the chambers were separated by a 8 µm filter, we recovered ∼ 1,000-fold less 100-23 bacteria in the presence of R2lc wild-type compared to the 100-23 monoculture (p < 0.001). In the presence of R2lc*Δpks*, levels of 100-23 were 3-fold less compared to the 100-23 monoculture (Figure 1E, left panels; p < 0.01). Viability of 100-23 was not impacted by R2lc when the chambers were separated by a 0.4 µm filter (Figure 1E, right panels). Since we did not observe antimicrobial activity when cultures were separated by a 0.4 µm membrane, which does allow flowthrough of metabolites, it is plausible that the antimicrobial compound is not secreted into the extracellular milieu.

To expand on these findings, we performed double-layer agar diffusion assays with a) filter-sterilized cell-free supernatant of R2lc, b) methanol and acetone extracts of the supernatant, c) live R2lc cells in MRS and PBS, d) R2lc cells killed following heat- or acetone treatment. Our findings further validate that the antimicrobial compound is not secreted (Figure S4AB); instead, only antimicrobial activity is observed when a cell suspension of viable R2lc (Figure S4C) is used whereas no zone of inhibition was observed with a cell suspension of viable R2lcΔ*pks* (Figure S4, bottom panel). Also, no zone of inhibition was observed when we inactivated R2lc by either heat or acetone treatment (Figure S4D). In conclusion, to elicit an antimicrobial effect, strain R2lc requires to be metabolically active and our data strongly suggest cell-cell interaction is required.

Next, we designed an experiment to determine if Pks-producing *L. reuteri* R2lc lyses the target cells using an analogous approach as we previously established (Alexander et al., 2019). We developed 100-23/leptin, a derivative of strain 100-23 that is engineered to accumulate leptin intracellularly. Following 12 hours batch culture of the 100-23/leptin monoculture, less than 15% of the total amount of recombinant leptin was released in the supernatant (Figure 1F, top panel). We expect that this relatively small amount of leptin is derived from natural lysis, for example, by biologically active prophages present in 100-23 (Oh et al., 2019). When R2lc Pks is co-cultured with 100-23/leptin, we found that a significant reduction of 100-23 CFU count directly correlates to an increase of leptin in the supernatant (Figure 1F, middle panel). However, the co-culture of R2lcΔ*pks* + 100-23/leptin (Figure 1F, bottom panel) yielded levels of 100-23 and leptin that were comparable to those observed in the 100-23/leptin monoculture (Figure 1F, top panel). Collectively, these data show that *L. reuteri* R2lc needs to be alive and likely requires cell-cell contact to elicit a Pks-mediated bactericidal effect that leads to lysis of the competitor strain.

### *L. reuteri* Pks molecules provide a competitive advantage *in vivo*

To characterize the ecological role of the *pks* antibacterial secondary metabolite gene cluster *in vivo*, we gavaged germ-free mice (n=4/group) with a 1:1 mixture of R2lc+100-23, or R2lcΔ*pks*+100-23. To quantify each strain in the fecal material, we engineered R2lc and its derivatives to encode chloramphenicol resistance (R2lc::*Cm* and R2lcΔ*pks::Cm*) while strain 100-23 was rifampicin-resistant (100-23—Rif^R^). We determined the competition ratio between R2lc and 100-23 (competition ratio: R2lc CFU count/competitor strain’s CFU count) over a period of six days in feces. We found that *L. reuteri* R2lc wild-type but not the Δ*pks* mutant gradually outcompetes the 100-23 strain. The R2lc:100-23 competition ratio increased from 107:1 (day 1) up to 980:1 (day 6), while the competition ratio between R2lcΔ*pks*:100-23 declined from 35:1 (day 1) to 4:1 (day 6) (Figure 2A). Similarly, the R2lc:100-23 competition ratio was higher than the R2lcΔ*pks*:100-23 competition ratio in the forestomach (31,973-fold vs 7.8-fold, *p* = 0.03), cecum (1,515-fold vs. 2.9-fold, *p* = 0.03), colon (1,474-fold vs. 8.9-fold, *p* = 0.03) and jejunum (1,114-fold vs. 5.2-fold, *p* = 0.03) (Figure 2B). Thus, our findings demonstrate that Pks provides *L. reuteri* R2lc with a competitive advantage throughout the murine intestinal tract.

**Figure 2.**
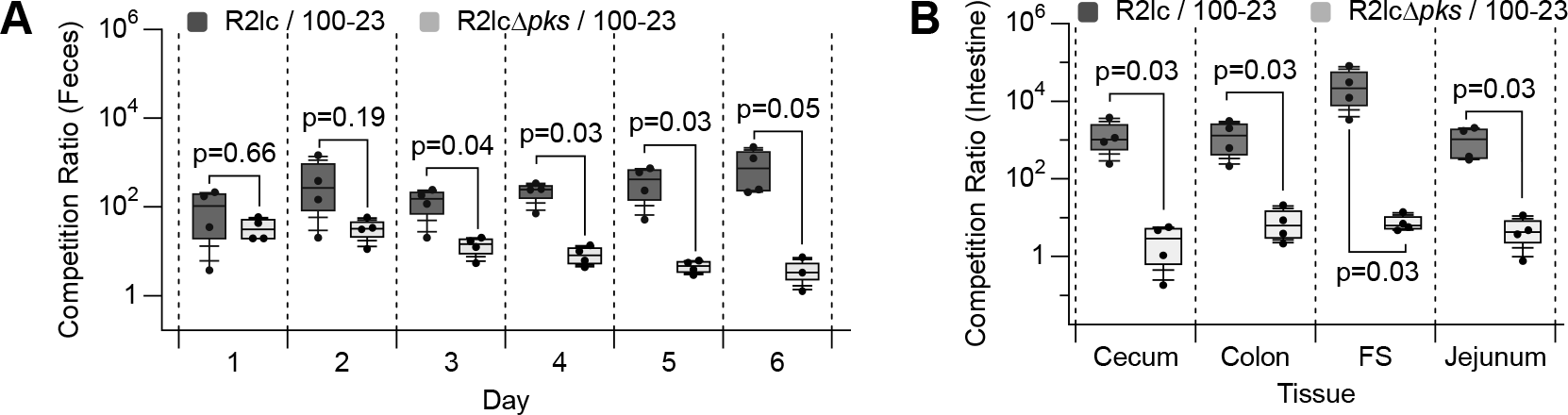
*L. reuteri* Pks molecules provide a competitive advantage *in vivo*. A 1:1 mixture of R2lc+100-23, or R2lcΔ*pks*+100-23 was administered by oral gavage to germ-free mice. **A)** Competition ratios of the indicated strains in fecal and **B)** intestinal content (day 6) were determined. In box and whisker plots, the whiskers represent the maximum and minimal values, and the lower, middle and upper line of the box represent first quartile, median and third quartile, respectively. Circles represent data from individual mouse. Statistical significance was determined by Wilcoxon / Kruskal-Wallis Tests; p<0.05 is considered statistically significant. F.S.: Forestomach.

### *L. reuteri* Pks-mediated antimicrobial effect is strain specific

Now we established that *L. reuteri* Pks provides an *in vivo* and *in vitro* competitive advantage against strain 100-23, we next investigated to what extent Pks-mediated killing is strain specific. We performed *in vitro* competition experiments with 51 *L. reuteri* gut isolates from different host origins: human, rat, mouse, chicken and pig. To quantify strains, we used derivatives of R2lc and R2lcΔ*pks* that were engineered to be chloramphenicol resistant. For each competitor strain we isolated a natural rifampicin-resistant mutant. Strain R2lc and the competitor strain were mixed 1:1 in MRS medium adjusted to pH 4.0, which is pH of the forestomach, the natural habitat of strain R2lc. The co-cultures were incubated for 24 hours, and appropriate dilutions were plated on MRS plates containing chloramphenicol (5 µg/ml, for R2lc and R2lcΔ*pks*) or rifampicin (25 µg/ml, for competitor strain). Competition ratios were determined after 24 hours of incubation.

Based on non-parametric statistical tests, we identified that R2lc inhibits 41 out of 51 (80.4%) *L. reuteri* strains. Specifically, six out of nine (66.6%) rat-isolates, seven out of ten (70%) mouse isolates, 16 out of 18 (89%) pig isolates, eight out of nine (88.9%) chicken isolates and two out of five (40%) human isolates were significantly inhibited by R2lc (R2lc vs competitor CFU, *P <0.05*). To test if the competitive advantage of R2lc is driven by Pks, we repeated the competition experiments with R2lcΔ*pks*. We found that deletion of the *pks* cluster resulted in loss of antimicrobial activity and the *Δpks* mutant did not show strong inhibition against competitor strains. However, Δ*pks* mutant still inhibited some competitor strains (16 out of 51) to a small extent, with observed differences in Colony Forming Units (CFUs) between Δ*pks* and competitor strains less than two orders of magnitude (Figure 3). These data demonstrate that R2lc-Pks mediated antimicrobial effect is strain specific and 10 out of 51 tested *L. reuteri* strains (19.6%) are not inhibited/outcompeted by the Pks producing strain.

**Figure 3.**
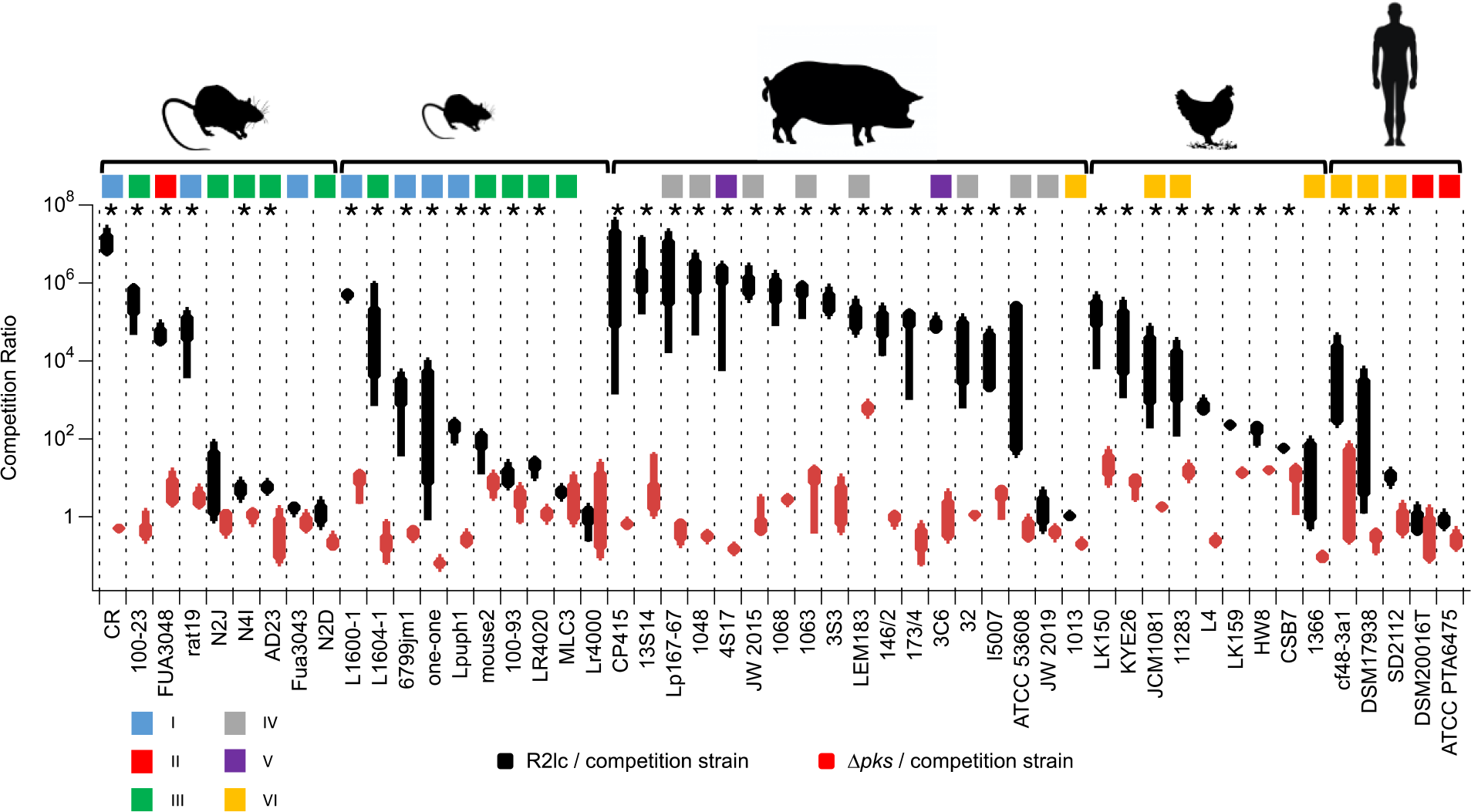
*In vitro* competition ratio of *L. reuteri* R2lc and R2lcΔ*pks* with a panel of 51 *L. reuteri* strains from different host origin. Competition ratios between R2lc and the competitor strain (black) and R2lcΔ*pks* and the competitor strain (red) after 24h co-incubation in MRS (pH 4.0). Data shown are based on at least three biological replicates. The whiskers represent the maximum and minimal values, and the lower, middle and upper line of the box represent first quartile, median and third quartile, respectively. Host origin (top) and the *L. reuteri* phylogenetic lineage (color coding) are indicated. Statistical significance was determined by Wilcoxon / Kruskal-Wallis Tests. Asterisks (*) represents statistical significance between R2lc vs competitor strain for the top panel (p<0.05 considered as significant). See also Table S3.

### *L. reuteri* strains resistant to Pks encode an acyltransferase enzyme

Next, we aimed to understand what the underlying mechanism is that allows select strains to be resistant to *L. reuteri* R2lc Pks. We performed comparative genome analyses to identify genes unique to strains that are resistant to *L. reuteri* R2lc Pks. We included 24 genome sequences in our analyses, of which four genome sequences were derived from resistant strains (mlc3, Lr4000, 6475 and 20016^T^) (Table S1). Our initial genome comparison analyses revealed seven genes unique to three of the four R2lc-resistant strains (mlc3, 6475 and 20016^T^) (Table S2).

One of the seven unique genes (*act*, Lreu_1368 in strain DSM20016^T^, Table S2) is annotated as an O-acyltransferase. This gene became our focus as several studies demonstrated that deletion of homologous *act* genes reduces the resistance to enzymes that target the peptidoglycan cell wall in other Gram-positive bacteria (reviewed in (Ragland and Criss, 2017)). Moreover, resistance to antimicrobial molecules is typically associated with altered cell-wall peptidoglycan structures (Vollmer, 2008). In *L. reuteri*, the *act* gene is located in the surface polysaccharide (SPS) gene cluster. Based on the available genome sequences, we found that three resistant strains (mlc3, 6475 and 20016^T^) all putatively encode a nearly identical Act protein (≥99% amino acid identity). Three R2lc-Pks*-*sensitive strains (CR, ATCC 53608 and one-one) putatively encode a distinct Act protein with 54% amino acid identity to the predicted Act amino acid sequence of resistant *L. reuteri* strains. Six additional R2lc-Pks*-*sensitive strains (I5007, 6799jm-1, lpup, CF48-3A1, Lr4020 and SD2112) putatively encode Act with 26-28% amino acid identity compared to the predicted Act amino acid sequence of resistant *L. reuteri* strains. The remainder of the 12 strains that are sensitive to R2lc-Pks do not contain the *act* gene in the SPS gene cluster (Figure 4).

**Figure 4.**
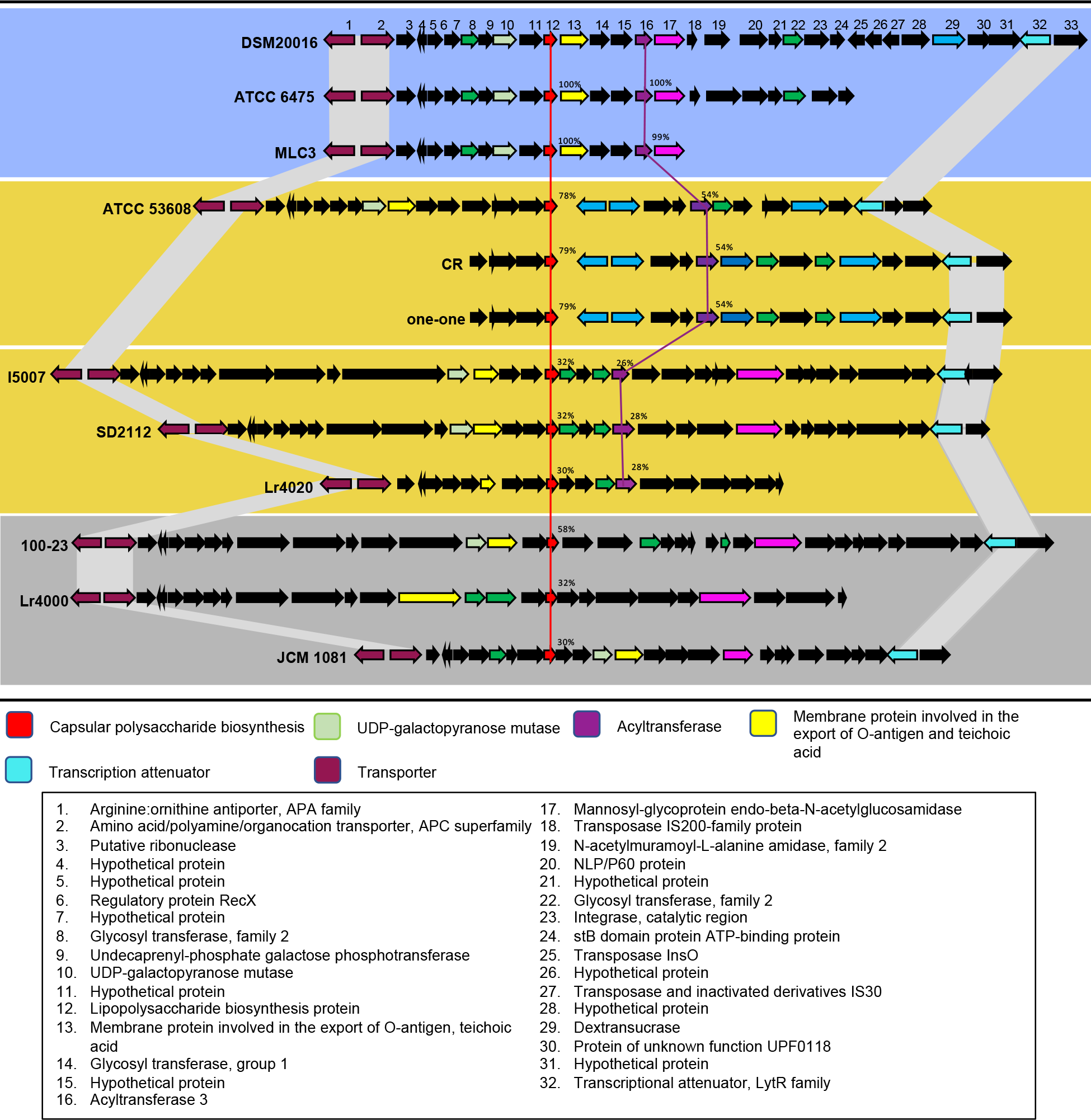
Variability in gene content of Surface Polysaccharide gene cluster in *L. reuteri.* Comparison of surface polysaccharide (SPS) gene clusters of strains resistant and sensitive to R2lc-Pks. Strains #1-3 (resistant) each putatively encode a nearly identical Act protein with ≥99% amino acid identity; strains #4-6 (sensitive) putatively encode Act that shares 54% amino acid identity to Act of resistant strains; strains #7-9 (sensitive) putatively encode Act that shares % amino acid identity to Act of resistant strains; strains #10-12 (sensitive) lack the *act* gene. Genes are color-coded with their predicted functions based on the annotation of the SPS gene cluster in *L. reuteri* DSM20016^T^. See also Table S1 and Table S2.

To map which *L. reuteri* strains are resistant or sensitive to R2lc-Pks, we constructed a phylogenetic tree of Act amino acid sequences from 134 *L. reuteri* strains. We found that 43 out of 134 strains (32.3%) contain an *act* gene whose putative product shares 99-100% amino acid identity to the predicted amino acid sequence of resistant *L. reuteri* strains, while the remaining strains putatively encode Act with 23-72% amino acid identity (Figure 5). Also see Table S6.

**Figure 5.**
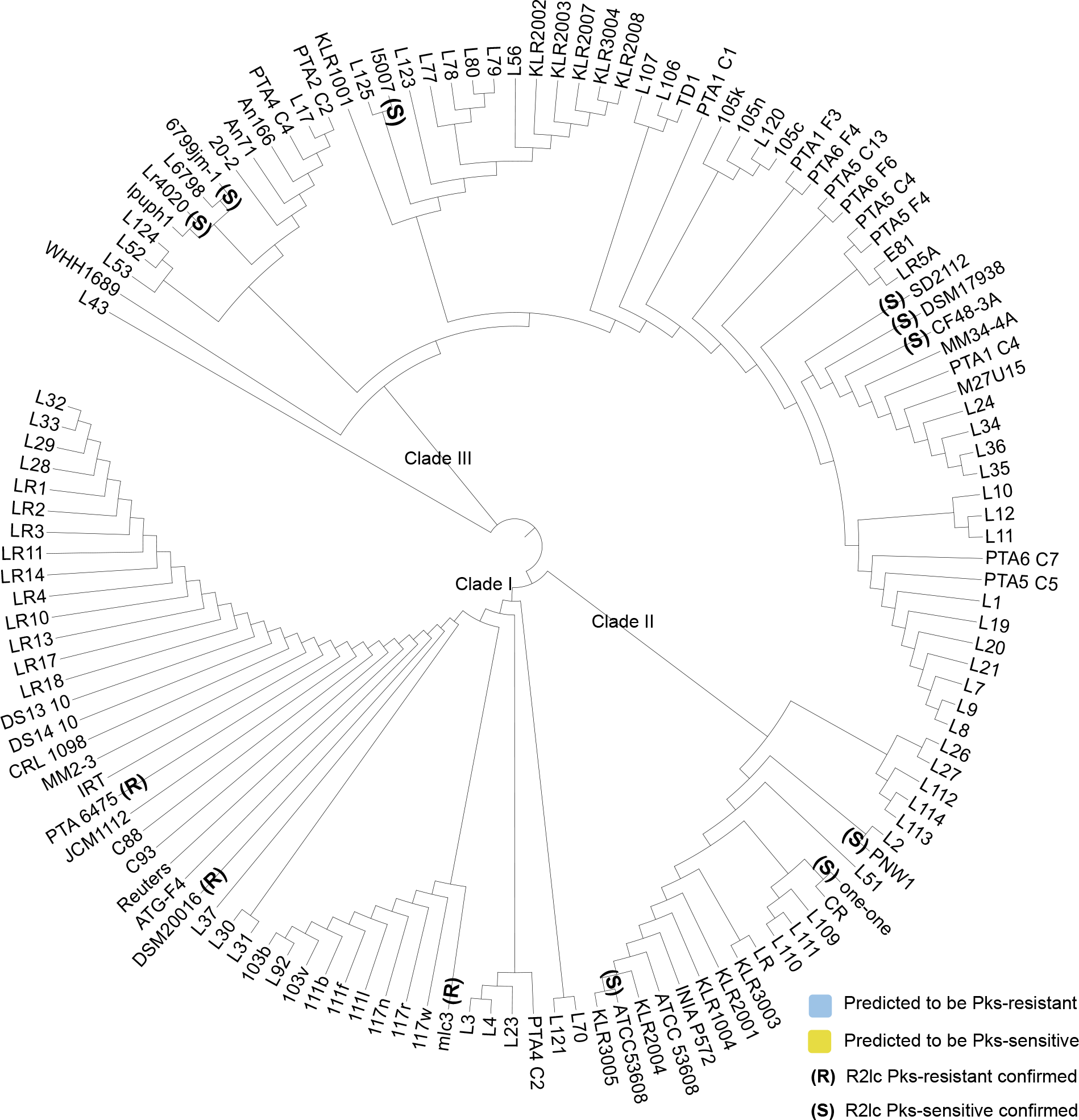
Phylogenetic tree of *L. reuteri* Act homologs. We constructed a phylogenetic tree based on Act amino acid sequences derived from 134 Act encoding *L. reuteri* strains to predict R2lc-resistance and sensitive strains. Forty-three out of 133 strains (32.3%) encode and *act* gene that shares 99-100% amino acid identify with the *act* gene in ATCC 6475 strain, thus, predicted to be R2lc-resistant (blue ring). The remaining strains encode an *act*gene that shares 23-54% amino acid identify with the *act* gene in PTA 6475 (yellow ring). See also Table S6.

### The acyltransferase gene in *L. reuteri* confers protection to R2lc-Pks products

To test if the *act* gene confers protection to R2lc-Pks products, we generated *act* mutants and complemented strains of mlc3 (rat isolate, clade I) and ATCC 6475 (human isolate, clade I). We observed that strain 6475Δ*act* but not mlc3Δ*act* grows slower *in vitro* compared to the parent wild-type strains. Growth was partially restored when 6475Δ*act* is complemented (with pNZ- *act*), while strain mlc3Δ*act* (with pNZ-*act*) grew faster than wild-type (Figure S5A). In these strains, we found that inactivation of *act* increased the sensitivity to R2lc-Pks, and *act* complementation decreased the sensitivity to R2lc-Pks, as is evident from the increased competition ratios between R2lc *vs* Δ*act* compared to the competition ratios between R2lc *vs* wild-type and complemented strain (Figure 6A, left panel). In contrast, deletion or complementation of *act* gene in mlc3 and 6475 did not change the competition ratio of these strains against R2lcΔ*pks* (Figure 6A, right panel). Overall, our data showed that Act in *L. reuteri* provides resistance to R2lc-Pks mediated killing.

**Figure 6.**
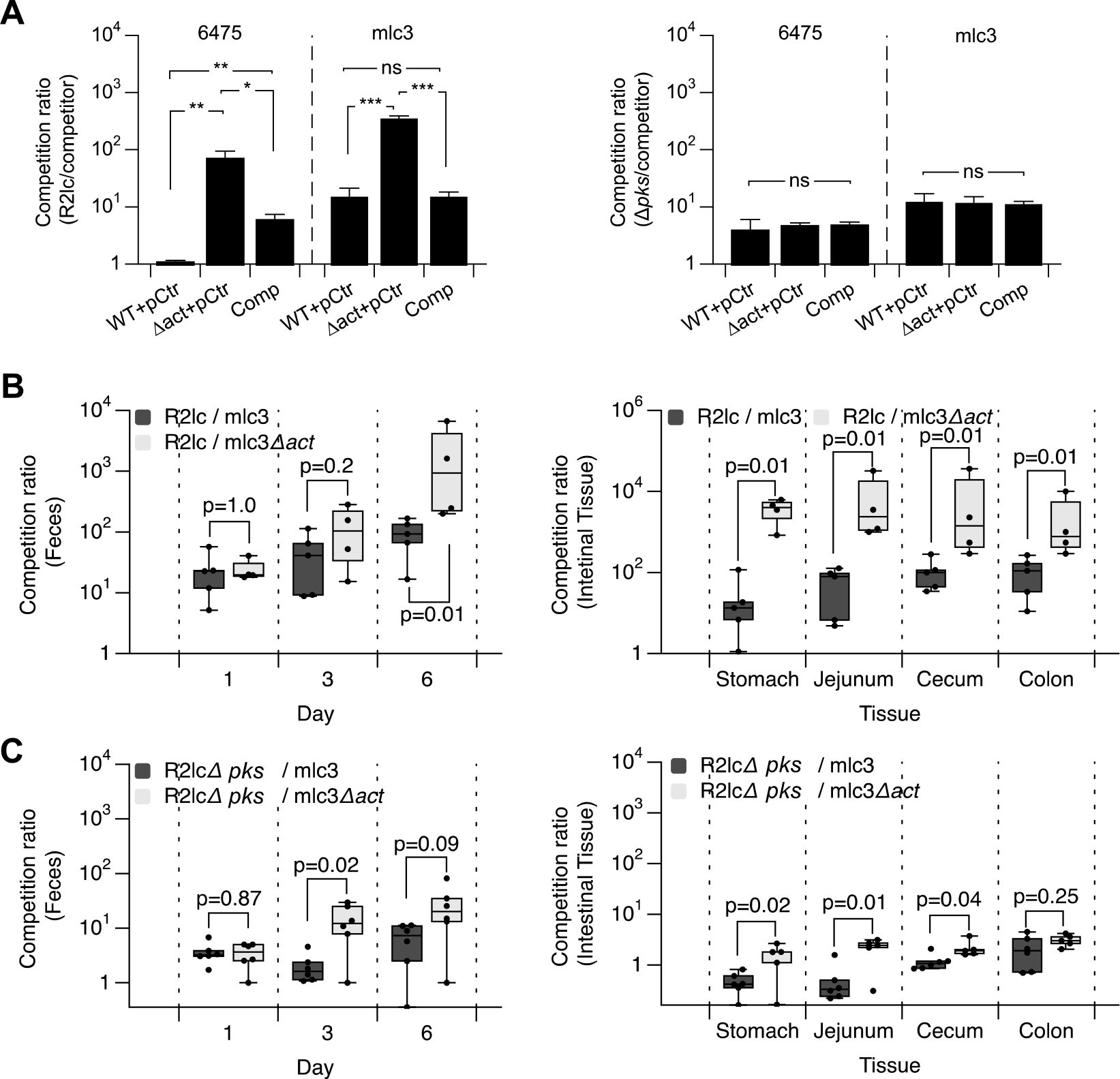
The *act* acyltransferase gene protects against R2lc-Pks molecules. **A)** Act provides resistance to R2lc-mediated killing (left panel), which is mediated by the inhibitory activity of an intact Pks (right panel). **B and C)** Competition ratios of the indicated strains in the previous germ-free mice in fecal (left panel) and intestinal contents (right panels). When data are presented as box and whisker plots, the whiskers represent the maximum and minimal values, and the lower, middle and upper line of the box represent first quartile, median and third quartile, respectively. Points represent individual data points. Statistical significance was determined by Wilcoxon / Kruskal-Wallis Tests, p=P value. P<0.05 considered significant. See also Figure S5.

### *L. reuteri* Act provides resistance against antimicrobial Pks molecules *in vivo*

To determine to what extent Act protects against R2lc Pks *in vivo*, we performed competition experiments in mice. Germ-free mice were administrated with a 1:1 mixture of any of the following strain combinations: R2lc ± *pks* and mlc3 ± *act* (Figure 6BC). Following administration, fecal samples were analyzed at day one, three and six, and animals were sacrificed at day 7 after which we analyzed microbial counts in the stomach, jejunum, cecum and colon. Fecal analyses revealed that the competition ratios between R2lc and mlc3 or its isogenic Δ*act* mutant were comparable (*P* > 0.05); however, at day 7, the competition ratio of R2lc:mlc3Δ*act* was significantly larger compared to R2lc:mlc3 (2,183 ± 3,060-fold *vs* 95 ± 58-fold, respectively; *P <* 0.01) (Figure 6B, left panel). We also observed that in all intestinal tissues the competition ratio of R2lc∷mlc3Δ*act* was larger compared to R2lc∷mlc3. Specifically, forestomach (3,777 ± 2,259-fold *vs* 31 ± 48-fold; *P* < 0.01), jejunum (9,510 ± 15,239-fold *vs* 62 ± 55-fold; *P* < 0.01), cecum (9,872 ± 17,683-fold *vs* 115 ± 99-fold; *P* < 0.01), colon (959 ± 4,703-fold *vs* 118 ± 104-fold; *P* < 0.01) (Figure 6B, right panel). Thus, the *act* gene provides protection against R2lc-Pks *in vivo*. To test to what extent R2lc Pks provides R2lc with a competitive advantage in these experiments, we performed competition experiments with R2lcΔ*pks*∷mlc3 and R2lcΔ*pks*∷mlc3Δ*act* (Figure 6C). Overall, the competition ratios of R2lcΔ*pks∷*mlc3 and R2lcΔ*pks∷*mlc3Δ*act* are lower compared to the competition ratios observed with R2lc wild-type, which demonstrates a clear role for Pks in providing R2lc with a competitive advantage *in vivo.* Interestingly, the R2lcΔ*pks*∷mlc3Δ*act* competition ratio is higher than that of R2lcΔ*pks*∷mlc3 in the stomach (1.5-fold vs 0.5-fold, *p* = 0.02), jejunum (2.2-fold vs 0.5-fold, *p* = 0.01) and cecum (2.2-fold vs 1.2-fold, *p* = 0.04) (Figure 6C, right panel), which could be linked to the role of Act in lysozyme resistance during *in vivo* colonization (Bera et al., 2005). Indeed, we tested the lysozyme susceptibility of wild-type and *act* mutant strains and identified that the Δ*act* variants are more susceptible to lysozyme while the complemented strains survived at levels comparable to that of the wild-type strains (Figure S5B). Therefore, host lysozyme may also suppress intestinal survival of mlc3Δ*act* thereby providing R2lcΔ*pks* a competitive advantage. When we tested the susceptibility of the *act* mutants to ampicillin we determined that the *act* mutants of 6475 (Figure S5C) and mlc3 (Figure S5D) are more sensitive to ampicillin, while the phenotype of complemented strains was similar to the wild-type strain.

Because Act provides protection to lysozyme and ampicillin, which both attack the peptidoglycan (Teethaisong et al., 2014; Vanderkelen et al., 2011), we aimed to understand how Act modifies the peptidoglycan complex.

### The acyltransferase gene increases acetylation of the cell wall

The bacterial Act enzyme acetylates the C6 hydroxyl of *N-*acylmuramic acid in peptidoglycan (PG) (Moynihan and Clarke, 2010). To understand the role of *L. reuteri* Act in acetylation, we performed HPLC analyses to determine acetylation levels of the cell wall in Pks- resistant and -sensitive strains, relative to total cellular muramic acid (MurN) content. Analysis of cell wall extracts derived from *L. reuteri* 100-23, a Pks-sensitive strain which does not encode Act, revealed that 6.65 ± 1.19% was acetylated relative to total MurN. Next, we cloned the *act* gene from strain mlc3 in the inducible expression vector pSIP411, which was established in strain 100-23. Induced expression of *act* in 100-23 increased relative acetylation levels to 28.7 ± 9.0%. Inactivation of *act* in *L. reuteri* strains resistant to R2lc-Pks reduced total acetylation levels (from 70.1 ± 4.9% to 2.3 ± 0.3% for mlc3; and from 61.2 ± 9.1% to 21.3 ± 12.9% for 6475). Overexpression of the *act* gene in mlc3Δ*act* and 6475Δ*act* restored the relative acetylation levels (from 2.3 ± 0.3% to % to 51.8 ± 15.1% for mlc3Δ*act::act* ; and from 21.3 ± 12.9% to 50.9 ± 9.2% for 6475Δ*act::act*) (Table 1). Together these data suggests that *L. reuteri* Act is responsible for increased acetylation of the cell wall, which we demonstrated is linked to the resistance to the antimicrobial compound produced by R2lc.

**Table 1.** O-acetylation levels were determined by base-catalyzed hydrolysis and release of acetate. Percent O-acetylation is reported relative to muramic acid (MurN) content.

| Strain | % O-acetylation |
| --- | --- |
| 100-23 | $6.65 \pm 1.19$ |
| 100-23 + pSIP control | $3.46 \pm 0.07$ |
| 100-23::act | $28.70 \pm 9.01$ |
| *mlc3* | $70.12 \pm 4.85$ |
| mlc3Δact + pSIP control | 2.31 ± 0.29 |
| mlc3Δact::act | 51.76 ± 15.09 |
| 6475 | 61.21 ± 9.07 |
| 6475Δact + pSIP control | 21.29 ± 12.92 |
| 6475Δact::act | 50.90 ± 9.18 |

<sup>a</sup> Results are from one representative biological replicate measured in triplicate and reported as mean with Standard Deviation.

## Discussion

In this study, we provided mechanistic insight into intra-species antagonism based on a secondary metabolite in the gut symbiont *L. reuteri*. We discovered that a biosynthetic gene cluster (BGC) provides a competitive advantage in the gastrointestinal tract by killing closely related microbes in a strain-specific manner. Expression of the act gene by select strains of *L. reuteri* increased resistance against this killing activity.

Closely related strains that occupy the same niche are subject to fierce competition (Hibbing et al., 2010; Segura Munoz et al., 2020). While the co-existence of two closely related strains may be stable in the host, long-term evolution may lead to different ecological outcomes (Edwards et al., 2018). Either evolution leads to changes that form a more stable population— lineages are overall less competitive—or evolution results in increased fitness of one lineage.

The lineage with increased fitness will replace the competing lineage, or the competing lineage may evolve in a way that will allow it to persist (Le Gac et al., 2012). Our findings are interesting in light of these ecological theories. Specifically, four distinct lineages have thus far been identified among rodent *L. reuteri* isolates (Frese et al., 2011). Our competition analyses revealed that nearly all strains in lineage I are sensitive to R2lc-Pks products while most strains in lineage III are resistant. One potential explanation for the latter is the broad distribution of nearly identical *act* genes in *L. reuteri* strains from different host origin (Figure 5, blue ring), which suggests *act* exerts an evolutionary pressure that will increase the fitness in the gut ecosystem. The findings presented in this work, the observation that aryl polyene gene clusters are widely distributed throughout the host-associated bacteria (Cimermancic et al., 2014), and that *L. reuteri* 6475Δ*act* has reduced gastrointestinal survival (preliminary data) support the importance of Act in gut fitness and make it collectively less likely that *act* is acquired through a neutral process such as genetic drift.

The evolution of a vertebrate symbiont with its host might be reciprocal, resulting in co-evolution. The beneficial traits that a microbe exerts on its host may have been shaped by natural selection as they promote host fitness, which is critical for the microbe to thrive in its niche.

R2lc-Pks molecules activates the aryl-hydrocarbon receptor (AhR), a ligand activated transcription factor (Özçam et al., 2019). Activation of AhR has been shown to induce production of interleukin-22 (IL-22) (Veldhoen et al., 2008), which enhances the innate immune response by inducing antimicrobial peptides (i.e. Reg3-lectins) production from the mucosal layer (Vaishnava et al., 2011; Zheng et al., 2008). It is intriguing to speculate that Pks molecules provide R2lc with a host-mediated competitive advantage as Pks-mediated AhR activation leads to antimicrobial peptide production by the host. Collectively, this would support the coevolution between select microbes and their host.

This work contributes to our understanding of the ecological role of a gut symbiont-encoded secondary metabolite and provides novel insight into how closely related strains developed resistance against these antimicrobial molecules by acetylation of the cell-wall polysaccharide complex. It is plausible bacteria have evolved other mechanisms to survive exposure to Pks. In this work, we did encounter one strain—the rodent isolate Lr4000—that does not encode Act but was resistant to R2lc Pks molecules during the batch culture competition experiments and the mechanism of resistance remains to be elucidated.

With a number of genome editing tools available for *L. reuteri*, we are in a position to use genetic approaches to study microbe-microbe interactions in a gut symbiont. When these experiments are placed in the right ecological context, mechanistic insight can be provided on the formation and stability of microbial communities, which is expected to provide insight how microbial diversity is maintained within complex ecosystems. This knowledge can be leveraged to provide rational approaches to select probiotics and next-generation probiotics to ultimately promote animal and human health.

## ACKNOWLEDGEMENTS

We thank Siv Ahrné (Lund University, Sweden) for providing *L. reuteri* R2lc, N2J, N2D, and N4I, BioGaia AB (Stockholm, Sweden) for providing *L. reuteri* strains ATCC PTA 6475, and Joseph Skarlupka for technical assistance. We are grateful to the College of Agricultural Life Sciences (CALS) Statistical Consulting Lab for their assistance in the statistical analysis, and the University of Wisconsin Biotechnology Center DNA Sequencing Facility for their services. This work was supported by startup funds from the University of Wisconsin-Madison to J.P.V.P., the UW-Madison Food Research Institute, and the United States Department of Agriculture, National Institute of Food and Agriculture (USDA NIFA) Hatch award MSN185615 and grant no. 2018- 6717-27523. M.Ö. received financial support from the Turkish Ministry of National Education and from the Department of Food Science. M.Ö. and S.Z. were the recipients of the Robert H. and Carol L. Deibel Distinguished Graduate Fellowship in Probiotic Research, which was awarded by the Food Research Institute (UW-Madison). J.C. is supported by National Institutes of Health grant R01 AI153173. Gnotobiotic work was partly supported by the Office of the Vice-Chancellor for Research and Graduate Education at the University of Wisconsin–Madison, with funding from the Wisconsin Alumni Research Foundation.

## AUTHOR CONTRIBUTIONS

M.Ö.: conceptualization, methodology, formal analysis, investigation, writing–original draft, writing–review&editing, visualization; J-H.O.: methodology, formal analysis, investigation, visualization; R.T.: methodology, validation, formal analysis, investigation; D.A.: methodology, validation, formal analysis; S.Z.: formal analysis, investigation, visualization; T.A.: validation, formal analysis, investigation; M.H.: validation, investigation; S.R-R.: validation; F.L.: investigation; C.C.C.: validation; E.V.: methodology; F.E.R.: methodology, resources; J.C.: investigation; T.B.: resources, supervision; J.W.: resources, writing—original draft, supervision; J.P.V.P.: conceptualization, methodology, writing–original draft, writing–review&editing, supervision, project administration, funding acquisition

## DECLERATION OF INTEREST

JPVP received unrestricted funds from BioGaia, AB, a probiotic-producing company. JPVP is the founder of the consulting company Next-Gen Probiotics, LLC. JW has received grants and honoraria from several food and ingredient companies, including companies that produce probiotics. JW is a co-owner of Synbiotic Solutions, LLC, and is on the Scientific Advisory Board of Alimentary Health. M.Ö. was an employee of DuPont Nutrition and Biosciences. JC is a Scientific Advisor for Seed Health, Inc.

## STAR METHODS

### RESOURCE AVAILABILITY

#### Lead contact

Further information and requests for resources should be directed to and will be fulfilled by the Lead Contact, Jan-Peter van Pijkeren.

#### Material Availability

Requests for the plasmids and strains used in this study should be directed to and will be fulfilled by the Lead Contact, Jan-Peter van Pijkeren.

#### Data and Code Availability

Any additional information required to reanalyze the data reported in this paper is available from the lead contact upon request.

### KEY RESOURCE TABLE

| REAGENT or RESOURCE | SOURCE | IDENTIFIER |
| --- | --- | --- |
| **Bacterial strains** |  |  |
| See Table S3 for bacterial stains used in this study |  |  |
| **Chemicals** |  |  |
| Agar | Alfa | Cat#A10752 |
| Agarose | IBI | Cat# IB70042 |
| LB broth | Acumedia, Neogen | Cat# 7290A |
| MRS broth | BD | Cat# 288110 |
| M17 broth | BD | Cat# 218561 |
| Potassium Buffered Saline | Gibco | Cat# 14190-144 |
| 0.45-μm-pore-size cellulose acetate membrane | Sigma | Product# WHA69012504 |
| 0.4-μm-pore-size filter insert | Cell Treat | Part no 230607 |
| 8-μm-pore-size filter insert | Cell Treat | Part no 230611 |
| Acetic acid standard solution | Fisher Scientific | Cat# A38S-500 |
| Sulfuric acid | Fisher Scientific | Cat# A300-500 |
| Protease | Sigma | Product# P5147 |
| Magnesium sulfate | Fisher Scientific | Cat# M65-500 |
| RNase A | Thermo Scientific | Cat# EN0531 |
| DNase I | Invitrogen | Cat# 18068015 |
| α-amylase | Sigma | Cat# 10065 |
| Sodium chloride | Fisher Scientific | Cat# S271-500 |
| Tris-HCl | Calbiochem | Product# 648310-M |
| Sodium dodecyl sulfate | Sigma | Product# L4509 |
| 24-well plates | Fisher Scientific | Cat# 12556006 |
| Agarose | DOT Scientific Inc. | Cat# DS170042 |
| Pellet paint | VWR International | Cat# 9049-3 |
| Chow diet | LabDiet | Cat# 5201 |
| DpnI | Fisher Scientific | Cat# FERFD1704 |
| 0.22 μm filter, Polyvinylidene fluoride [PVDF] | Millipore | Cat# SLGVM33RS |
| Sodium hydroxide | Fisher Chemical | Cat# S318100 |
| Methanol | Fisher Chemical | Cat# A456-500 |
| Formic acid | Fisher Chemical | Cat# A11350 |
| Chloramphenicol | Dot Scientific | Cat# DSC61000-25 |
| Erythromycin | Fisher Scientific | Cat# BP920-25 |
| Rifampicin | TCI Chemicals | R0079-25G |
| Vancomycin | Chem-Impex | Cat# 00315 |
| Ampicillin | DOT Scientific | Cat#DSA40040-25 |
| Ethanol | KOPTEC | Cat# V1016 |
| P<sub>sak</sub> Induction peptide | Peptide2.0 | Sequence:<br>MAGNSSNFIHKIKQIFTHR |
| Choice Taq DNA polymerase master mix | Denville Scientific | Cat# CB4070-8 |
| Phusion Hot Start polymerase II | Thermo Scientific | Cat# F-549-L |
| T4 DNA ligase | Thermo Scientific | Cat# EL0011 |
| Polynucleotide kinase | Thermo Scientific | Cat# EK0031 |
| dATP | Promega | Cat# U120A |
| Ampligase | Lucigen | Cat# A32750 |
| Sodium Hydroxide | Fisher Scientific | Cat# S318-500 |
| HCl | Ricca Chemical Company | Cat# 3440-1 |
| Zirconia/silica beads | Bio Spec | Cat#11079101Z |
| **Critical Commercial Assay Kits** |  |  |
| Qubit dsDNA quantification kit | Invitrogen | Cat# Q32853 |
| Genomic DNA purification kit | Promega | Cat# A1120 |
| GeneJET PCR purification kit | Thermo Scientific | Cat# K0701 |
| Plasmid isolation kit | Promega | Cat# A9340 |
| Leptin ELISA kit | R&D systems | Cat# MOB00B |
| **Experimental Models:** |  |  |
| **Organisms/Strains** |  |  |
| C57BL/6J, Male | Jackson Laboratory | Cat# 000664 |
| **Oligonucleotides** |  |  |
| See Table S4 for primers used for this study |  |  |
| **Recombinant DNA** |  |  |
| See Table S5 for plasmids used for this study |  |  |
| **Software and Algorithms** |  |  |
| JMP Pro | SAS | Ver. 11.0.0 |
| Datagraph | Visual Data Tools Inc. | Ver. 4.2.1 |
| Bruker Hystar software | Bruker | Hystar 3.2. |
| MEGA | (Kumar et al., 2018) | Ver. 10.1.7 |

### EXPERIMENTAL MODEL AND SUBJECT DETAILS

#### Microbial Strains and Growth Conditions

The strains and plasmids used in this study are listed in Table S3 and S5. *L. reuteri* strains were cultured in De Man Rogosa Sharpe (MRS) medium (Difco, BD BioSciences). For growth curve experiments, *L. reuteri* strains were grown for ∼16 hours and transferred to pre-warmed MRS with a starting inoculum of OD600= 0.1. Where appropriate, antibiotics were added to the medium as described below. We determined the OD600 at regular intervals as shown in growth curve figures. For *in vitro* competition and biofilm formation experiments, we used filter-sterilized (0.22 µm PVDF filter, Millipore) MRS that was adjusted to pH 4.0 with 1 M HCl.

Unless stated otherwise, we prepared bacterial cultures as follows: *L. reuteri* strains were incubated at 37°C under hypoxic conditions (5% CO2, 2% O2). The MRS agar plates were incubated for 24 hours for colony counting. *Escherichia coli* EC1000 was used as general cloning host and cultured with aeration at 37°C in lysogeny broth (LB, Teknova). *Lactococcus lactis* MG1363 harboring pJP042 (VPL2047) was cultured under static condition at 30°C in M17 broth that was supplemented with glucose (0.5% [w/v]). Electrocompetent *E. coli* EC1000 were prepared as described in (Sambrook and Russell, 2006). Electrocompetent *L. reuteri* cells were prepared as described in (Van Pijkeren and Britton, 2012). If applicable, MRS for *L. reuteri* was supplemented with 5 µg/ml erythromycin, 5 µg/ml chloramphenicol or 25 µg/ml rifampicin.

### Mice

#### Ethics statement

All mouse experiments were performed in accordance with NIH guidelines, Animal Welfare Act, and US federal law and were approved by the Application Review for Research Oversight at Wisconsin (ARROW) committee and overseen by the Institutional Animal Care and Use Committee (IACUC) under protocol ID M005149-RO1-A01. Conventional pathogen-free and germ-free mice were housed at Animal Science and Laboratory of Animal Research Facilities respectively at the University of Wisconsin-Madison.

#### Mice Strains and Husbandry

Twelve-week-old germ-free male B6 mice (C57BL/6) were maintained in a controlled environment in plastic flexible film gnotobiotic isolators with a 12 hours light cycle. Sterilized food (standard chow, LabDiet 5001, St. Louis, MO) and water were provided *ad libitum*.

#### Reagents and Enzymes

To amplify DNA fragments for cloning and screening, we used Phusion Hot Start DNA Polymerase II (Thermo Scientific) and Taq DNA polymerase (Denville Scientific), respectively. We used T4 DNA ligase (Thermo Scientific) for blunt-end ligations. If applicable, we treated column purified (Thermo Scientific) PCR products with DpnI (Thermo Scientific) to remove plasmid template DNA. Phosphorylation of double stranded DNA fragments was performed with T4 Polynucleotide Kinase (Thermo Scientific). Ligase Cycling Reactions (LCR) were performed as described previously (Kok et al., 2014). Oligonucleotides and double-stranded DNA fragments were synthesized by Integrated DNA Technologies (IDT; Table S2).

### METHOD DETAILS

Construction of *L. reuteri* mutant strains

#### Construction of suicide shuttle vectors for homologous recombination

To generate mutant strains in lactobacilli we used the counterselection plasmid pVPL3002 (Zhang et al., 2018). To generate *L. reuteri* R2lc*∷Cm,* R2lcΔ*pks::Cm,* mlc3Δ*act* and DSM17938Δ*act*, 500-1000 bp upstream and downstream flanking regions of target genes were amplified by PCR (see Table S4 for oligonucleotides). We used following oligonucleotides; oVPL3113-3114 (upstream, R2lc::*Cm*), oVPL3115-3116 (downstream, R2lc::*Cm*), oVPL3194-3195 (upstream, mlc3Δ*act*), oVPL3196-3197 (downstream, mlc3Δ*act*), oVPL3237-oVPL3238 (upstream, DSM17938Δ*act*) and oVPL3239-oVPL3240 (downstream, DSM17938Δ*act*).

Amplicons were purified (GeneJET PCR Purification Kit, Thermo Scientific). The pVPL3002 backbone was amplified with oVPL187-oVPL188, purified (GeneJET PCR Purification Kit, Thermo Scientific) and digested with DpnI. Purified amplicons were quantified (Qubit, Life Technologies). The amplicons were mixed at equimolar quantities (0.25 pmol), followed by phosphorylation, ethanol precipitation and LCR (Kok et al., 2014). Following LCR, DNA was precipitated with Pellet Paint co-precipitant (VWR International), resuspended in 5 µl sterile water and transformed into electrocompetent *E. coli* EC1000 cells. By PCR, we screened for insertion of our target sequences using oligonucleotides that flank the multiple cloning site (oVPL49-oVPL97). Finally, the integrity of deletion and insertion cassettes was determined by Sanger sequencing. The resultant constructs were named as follow; VPL31130 (contains the chloramphenicol insertion cassette for R2lc::*Cm* and R2lcΔ*pks::Cm*), and VPL31079 (contains *act* deletion cassette for mlc3).

#### Construction of *L. reuteri* mutants by homologous recombination

Three micrograms plasmid DNA was electroporated in electrocompetent *L. reuteri* cells. For R2lc::*Cm,* bacterial cells were plated on MRS agar containing 5 µg/ml chloramphenicol, colonies were screened by PCR (oVPLl3117-3118) to confirm double crossover event. For mlc3Δ*act,* bacterial cells were plated on MRS agar containing 5 µg/ml erythromycin and colonies were screened for single crossover (SCO) event by PCR with oligonucleotide mixtures oVPL3216- oVPL3217-oVPL97 (upstream SCO) and oVPL3216-oVPL3217-oVPL49 (downstream SCO). Following confirmation of SCO, bacterial cells were cultured for one passage in MRS broth without antibiotic selection, and cells were plated on MRS agar plates containing 400 µg/ml vancomycin. Vancomycin-resistant colonies represent cells in which a second homologous recombination event took place (Zhang at al. 2018). To confirm double crossover (DCO), we performed PCR using oligonucleotides oVPL3216-oVPL3217 (for mlc3Δ*act*) and oVPL3244- oVPL3245 (for DSM17938Δ*act*). We used Sanger sequencing to verify the integrity of the recombinant strains.

#### Inactivation of *L. reuteri* 6475 acyltransferase by recombineering

To inactivate the gene putatively encoding acyltransferase (LAR_1287) in *L. reuteri* 6475, we applied single-stranded DNA recombineering using previously established procedure (Van Pijkeren et al., 2012). An 80-mer oligonucleotide identical to the lagging strand with five consecutive mismatches yielded, when incorporated, two in-frame stop codons (Van Pijkeren and Britton, 2012). Electrocompetent *L. reuteri* 6475 cells harboring pJP042 (VPL2047) were transformed with 100 µg oVPL3166. To identify recombinants, we plated cells on MRS plates and recombinant genotypes were identified by mismatch amplification mutation assay (MAMA) PCR (Cha et al., 1992) using oligonucleotide combination oVPL3163-3164-3165. After colony purification, the pure genotype recombinants were confirmed by MAMA-PCR and Sanger sequencing. The resulted mutant strain was named as *L. reuteri* 6475Δ*act.* To cure pJP042, cells were grown in plain MRS for two passages and plated on MRS agar. Then cells were inoculated in MRS containing 5 µg/ml erythromycin to identify cells from which pJP042 was cured.

#### Construction of Δ*act* mutants, pSIP controls, and complemented strains

To complement *act* into *L. reuteri* 6475Δ*act* (VPL4241) and mlc3Δ*act* (VPL4308), we amplified by PCR the *act* gene from *L. reuteri* 6475 and mlc3 (oVPL3188-3189), the EFtu promoter from *L. reuteri* 6475 (oVPL 1447-1448) and the pNZ8048 backbone (oVPL309-310). PCR products were quantified, purified (Fisher Scientific, GeneJET), and DpnI treated (Fisher Scientific, FastDigest). Subsequenlty, amplicons were purified again and mixed at equimolar concentrations (0.25 pmol), phosphorylated (T4 polynucleotide kinase, Thermo Scientific), and ethanol precipitated. DNA fragments were assembled by Ligase Cycling Reactions (LCR) (oVPL 4262-4264), followed by precipitation with Pellet Paint co-precipitant (VWR International), resuspension in 10 μL sterile water and transformation into electrocompetent EC1000 cells. By colony PCR inserts were identified (oVPL1447 and oVPL3188), and the integrity of the constructs was verified by Sanger sequencing. The resulting strains were labeled *E. coli* VPL31603 and VPL31601 which contain pNZ8048-*actmlc3* and pNZ8048-*act6475*, respectively. Hereafter, pNZ8048-*actmlc3* and pNZ8048-*act6475* are referred to as pVPL31603 and pVPL31601, respectively.

One microgram of pVPL31603 and pVPL31601 were transformed into electrocompetent rifampicin-resistant *L. reuteri* mlc3Δ*act* (VPL4248) and *L. reuteri* 6475Δ*act* (VPL4241), respectively. Cells were plated on MRS containing 5 μg/mL erythromycin and screened via colony PCR (oVPL1447 and oVPL3188). The mutant strains were labeled VPL31666 (mlc3Δ*act* + pVPL31603) and VPL31664 (6475Δ*act* + pVPL31601). Using the above method, control strains were generated by transforming pNZ8048 empty vector into rifampicin-resistant *L. reuteri* mlc3, mlc3Δ*act*, 6475, and 6475Δ*act*, and were labeled VPL 31668, 31675, 31613, 31673, respectively.

#### Generating rifampicin resistant *L. reuteri* strains

To enumerate the bacteria co-cultured with R2lc∷*cm*, we generated mutants that naturally acquired mutations to yield resistance to rifampicin. Briefly, *L. reuteri* strains were grown in MRS for 16 hours and plated on MRS agar containing 25 µg/ml rifampicin followed by 24 hours of incubation at 37°C under hypoxic condition. Rifampicin resistant colonies were purified by a conventional streak plate method and the corresponding cultures were stored at -80°C.

#### Whole Genome Sequencing

Genomic DNA was isolated using the genomic DNA purification kit (Wizard, Promega), and DNA concentrations were determined using the Qubit® (dsDNA High Sensitivity Assay Kit, Life Technologies). Whole genome sequencing was performed at the University of Wisconsin-Madison Biotechnology Center. Samples were prepared according to the Celero PCR Workflow with Enzymatic Fragmentation (NuGEN). Quality and quantity of the finished libraries were assessed using an Agilent bioanalyzer and Qubit^®^ dsDNA HS Assay Kit, respectively. Libraries were standardized to 2 nM. Paired end, 250 bp sequencing was performed using the Illumina NovaSeq6000. Images were analyzed using the standard Illumina Pipeline, version 1.8.2.

#### Comparative Genome and Bioinformatic Analyses

Reads were assembled *de novo* using SPAdes (version 3.11.1) software. Assembled draft genomes were uploaded to National Center of Biotechnology Information (NCBI) ((Özçam and van Pijkeren, 2019; Zhang et al., 2020)) and the Joint Genome Institute Integrated Microbial Genomes and Microbes (JGI-IMG) database to perform comparative genome analyses. The web-based comparative genome database Phylogenetic Profiler from the JGI-IMG was used to identify genes present only in *L. reuteri* strains that are resistant to R2lc (ATCC PTA6475 [named as MM4- 1A in JGI-IMG], mlc3, DSM20016 and Lr4000).

We used *L. reuteri* JCM112 acyltransferase protein sequence (LAR_1287) as a query sequence to search the National Center for Biotechnology Information (NCBI) Nonredundant protein sequence (nr) and JGI-IMG database to identify homologous among *L. reuteri* strains. Partial protein sequences were excluded from the data set. Amino acid sequences of *L. reuteri act* genes were aligned by using MUSCLE (Edgar, 2004), and we constructed the phylogenetic tree with MEGA 7.0 software (Kumar et al. 2016).

#### *In vitro* competition assay with *L. reuteri* strains

*L. reuteri* strains were grown in MRS for 16 hours. The rifampicin resistant competition strains were co-cultured with either R2lc::*Cm* or R2lcΔ*pks::Cm* at optical density OD600: 0.05 (from each strain in co-culture) in a pre-warmed MRS broth (pH 4.0). The co-cultures were then incubated for 24 hours at 37°C. After incubation, a serial dilution was performed and 100 µl from appropriate dilution was plated to MRS plates containing chloramphenicol (5 µg/ml) or rifampicin (25 µg/ml). After 24 hours of incubation, we determined the competition ratios.

#### *In vitro* biofilm assay

*L. reuteri* R2lc, R2lcΔ*pks* and 100-23 cultures were inoculated into 2 ml MRS broth at OD600= 0.1 (0.05 from each strain if co-culture) in 24-well plates (Fisher Scientific) and incubated for 24 hours. The culture supernatant was removed, and bacterial biofilm was washed three times with 1 ml PBS. The adherent cells were scraped from the well and re-suspended in 2 ml PBS. For colony enumeration, the suspended cultures were plated on MRS plates containing chloramphenicol (5 µg/ml) for R2lc and R2lcΔ*pks* or rifampicin (25 µg/ml) for 100-23, and incubated for 24 hours.

#### Split-well (filter separated) competition experiment

Two cultures (R2lc:Cm^R^ or R2lc *Δpks*:Cm^R^ vs. 100-23, Rif^R^) were separated by two inserts with different membrane pore sizes in 6-well plates as shown in Figure 1E. Filter inserts (0.4 or 8 µm; CellTreat) were placed in a 6-well plate to create upper and lower layers.

Overnight cultures (16 hours at 37°C in MRS) of 100-23 and either R2lc:Cm^R^ or R2lc*Δpks*:Cm^R^ were washed once in MRS (pH= 4.0) by centrifugation (5 min at 5,000 × *g*), resuspended in MRS to make initial OD600= 0.05. Three mL of R2lc:Cm^R^ (or R2lc *Δpks*:Cm^R^) and 100-23 (Rif^R^) suspensions were added to upper (R2lc:Cm^R^ or R2lc*Δpks*:Cm^R^) and lower wells (100-23 [Rif^R^]), respectively. Plates were incubated for 24 hours at 37°C under hypoxic conditions. The viability of the cells (CFU/mL) was determined by the standard plate count method as described above.

#### Sample Preparation for Double-Layer Agar Antimicrobial Assay

Live cells from R2lc:Cm^R^ or R2lc *Δpks*:Cm^R^ overnight cultures (16 hours) were prepared in MRS or PBS following a wash with MRS and PBS, respectively. Bacterial culture supernatant was collected following centrifugation (1 min at 15,000 × *g*). Heat-treated (dead) cells were prepared following heat treatment at 70°C for 30 min. For secondary metabolite extraction from cells, acetone (1 vol) or methanol (1/50 vol) was added to the washed cell pellets obtained from fresh culture followed by shaking at 500 rpm at room temperature for 3 hours (Eppendorf, Thermomixer 5382). Before drop plating, all supernatant and cell extracts were filter-sterilized (0.22 µm, EMD Millipore).

#### Double-Layer Agar Antimicrobial Assay

Bacterial culture, supernatant, and cell extracts of R2lc:Cm^R^ or R2lc *Δpks*:Cm^R^ were subjected to the antimicrobial assay using the double-layer agar containing 100-23 cells. Two hundred μL of 100-23 overnight culture diluted in MRS (OD600= 0.5) was mixed with 3 mL of pre-warmed (50°C) 0.2% agarose in dH2O followed by pouring the mixture onto the MRS-agar plate. Solidified top-agar was dried for 15 min at room temperature. Ten μL of each sample was dropped onto the top agar and dried until completely absorbed. All double-layer agar plates were incubated at 37°C under a hypoxic condition (2% O2, 5% CO2) for 24 hours.

#### Construction of leptin producing 100-23 strain and leptin release experiment

Rifampicin-resistant 100-23 strain (VPL4251) was transformed with pVPL31131, the plasmid that encodes murine leptin (Alexander et al., 2019), to yield strain VPL31598 (100- 23/leptin). Cm^R^-derivatives of *L. reuteri* R2lc (VPL4231) and R2lcΔ*pks* (VPL4209), and 100- 23/leptin were grown in MRS, the latter supplemented with 5 µg/ml erythromycin. One ml of each bacterial culture was centrifuged (2 min 16,000 × *g*) and washed in an equal volume of MRS, which was subsequently added to 40 mL pre-warmed MRS. Mono-culture experiments for 100-23/leptin and co-culture experiments for R2lc + 100-23/leptin, R2lcΔ*pks* +100-23/leptin were initiated with a starting inoculum of OD600= 0.1. At 0, 2, 4, 6, 8, and 10 hour time points, we collected samples of each culture and plated on MRS-agar supplemented with 25 µg/ml rifampicin to determine 100-23/leptin CFU counts. In addition, we collected samples of each culture to harvest cell-free supernatant and total cell lysate for leptin quantification. Nine hundred microliters culture were centrifuged for 1 minute at 16,000 × *g*, filter-sterilized, and stored at -20°C. To obtain total cell lysate, we used a bead beater (BioSpec) to lyse 1 mL culture samples in tubes containing ∼100 µL 0.1 mm zirconia/silica beads with 3 minutes total bead beating time, performed in two 90 seconds stages with 1 min cooling on ice in between. After bead beating, samples were spun down (1 minute at 10,000 rpm) and the supernatant was filter-sterilized and stored at -20°C. Supernatant and cell lysate samples were thawed on ice and leptin levels were quantified by ELISA.

#### *In vivo* competition experiment

Germ-free B6 (C57BL/6J, male 12-week-old, n=4-5mice/group) mice were maintained in sterile biocontainment cages in the gnotobiotic animal facility at the University of Wisconsin-Madison. Mice were colonized following a single oral gavage of 200 μl *L. reuteri* cocktail in PBS (1:1 ratio, ∼2×10^8^ CFU). Each group was gavaged with a mixture of either *L. reuteri* R2lc::*Cm* + competition strain or *L. reuteri* R2lcΔ*pks::Cm* + competition strain. Twenty-four hours following colonization, fecal samples were collected daily from individual mice to determine the fecal CFUs. Fecal samples were homogenized and diluted in PBS followed by plating on MRS-Cm and MRS-Rif. After 24 hours of incubation colonies were counted. At day 7, mice were sacrificed by CO2 asphyxiation and digesta from forestomach, jejunum, cecum and colon were collected, weighed and re-suspended with PBS (100 mg/ml) to determine the CFUs per 100 mg content.

#### Ampicillin Resistance Experiment

*L. reuteri* 6475 + pCtl, 6475Δ*act* + pCtl, 6475Δ*act +* pNZ8048-*act6475*, mlc3 + pCtl, mlc3Δ*act* +pCtl, and mlc3Δ*act +* pNZ8048-*actmlc3*, were grown in MRS supplemented with 5 µg/ml erythromycin for 16 hours. The cultures were centrifuged, washed twice in 1 mL MRS, and sub-cultured into 10 mL pre-warmed MRS with no antibiotic. Sub-cultures grew to OD600=0.6 and subsequently sub-cultured again into 1 mL aliquots containing a range of concentrations of ampicillin with a starting inoculum of OD600=0.01. Samples were transferred to a 96-well flat-bottom plate for 24 hour growth curve measurements using a MultiSkan Sky Plate Reader (ThermoFisher, Waltman, MA, USA) with 37°C, kinetic loop with 5 second shaking intervals every 30 min.

#### Determination of peptidoglycan O’-acylation by HPLC

Peptidoglycan (PG) was extracted following a previously reported method (Ha et al., 2016) with some modifications. After 16 hours of incubation, the cells were harvested by centrifugation at 5,000 × *g* and 4 °C for 5 min, washed twice with 10 mM sodium phosphate buffer (pH 6.5) and then resuspended in 50 ml of water (pH 5.5 to 6.0). The cell suspension was boiled in an equal volume of 8% (w/v) sodium dodecyl sulfate (SDS, 4% w/v final concentration) in 25 mM SP buffer (pH 6.5) for 1 hour under reflux with stirring. Samples were centrifuged (30,000 × *g*) and the SDS-insoluble PG was washed (5 times) with sterile double distilled H2O to completely remove SDS and lyophilized. Lyophilized PG was dissolved in 4 ml of 1:1 mixture of 10 mM Tris-HCl (pH 6.5) and 10 mM NaCl and sonicated for 2 min. The PG suspension was treated with 100 µg/ml α-amylase (from *Bacillus sp*., Sigma), 10 µg/ml DNase I (Invitrogen), 50 µg/ml RNase A (Thermo Scientific), and 20 mM MgSO4 solution and incubated overnight at 37 °C with shaking. The PG suspension was further treated with 200 µg/ml protease (from *Streptomyces griseus*, Sigma) and incubated overnight at 37 °C with shaking. Samples were then re-extracted by boiling in 1% SDS for 40 min, washed, lyophilized and stored at -20 °C until use.

For the analysis of acetate release from PG, lyophilized PG (20 mg) was dissolved in 150 µl ddH2O and mixed with an equal volume of either 160 mM NaOH or 160 mM sodium phosphate buffer (pH 6.5) and incubated overnight at 37°C with shaking. The peptidoglycan was collected by centrifugation at 15,000 × *g* for 20 min. The supernatant was filtered through a 0.45-µm-pore-size cellulose acetate membrane and quantitation of released acetate was carried out in a Dionex UltiMate 3000 HPLC equipped with an LPG-3400 quaternary pump, a WPS-3000 analytical autosampler, and a DAD-3000 diode array detector. The filtered supernatants or 100 µM of acetic acid standard solution were injected onto a 300 × 7.8 mm Rezex^TM^ ROA-Organic Acid H^+^ (8%) column and eluted isocratically with 20 mM H2SO4 at 0.5 ml min^-1^. Total MurNAc was liberated by acid-catalyzed hydrolysis (6 N HCl, 100°C for 1.5 hour). MurNac was then determined by HPLC and quantified against a calibration curve established using standard MurNac. The column was maintained and 40 °C and absorbance was monitored at 210 nm.

#### LC/MS Sample preparation

An aliquot of 1.5 ml from each of the cultures were collected in Eppendorf tubes and centrifuged at 2152 × *g* for 5 min. The supernatants were collected and transferred to dram vials and the cell pellets were incubated for 1 hour in 100 μl methanol. Next, the samples were centrifuged again and the methanol extracts were added to 1 dram vials. A Gilson GX-271 liquid handling system was used to subject 900 μL of the samples to automated solid phase extraction (SPE). Extracts were loaded onto pre-conditioned (1 ml MeOH followed by 1 ml H2O) EVOLUTE ABN SPE cartridges (25 mg absorbent mass, 1 ml reservoir volume; Biotage, S4 Charlotte, NC).

Samples were subsequently washed using H2O (1 ml) to remove media components, and eluted with MeOH (500 μL) directly into an LC/MS-certified vial.

#### UHPLC/HRESI-qTOF-MS Analysis of Extracts

LC/MS data were acquired using a Bruker MaXis ESI-qTOF mass spectrometer (Bruker, Billerica, MA) coupled with a Waters Acquity UPLC system (Waters, Milford, MA) with a PDA detector operated by Bruker Hystar software, as previously described (Hou et al., 2012). Briefly, a solvent system of MeOH and H2O (containing 0.1% formic acid) was used on an RP C-18 column (Phenomenex Kinetex 2.6μm, 2.1 × 100 mm; Phenomenex, Torrance, CA) at a flow rate of 0.3 ml/min. The chromatogram method started with a linear gradient from MeOH/ H2O (10%/90%) to MeOH/ H2O (97%/3%) in 12 min, then held for 2 min at MeOH/ H2O (97%/3%). Full scan mass spectra (m/z-150-1550) were measured in positive Electrospray Ionization (ESI) mode. The mass spectrometer was operated using the following parameters: capillary, 4.5 kV; nebulizer pressure,

1.2 bar; dry gas flow, 8.0 L/min; dry gas temperature, 205 °C; scan rate, 2 Hz. Tune mix (ESI-L low concentration; Agilent, Santa Clara, CA) was introduced through a divert valve at the end of each chromatographic run for automated internal calibration. Bruker Data Analysis 4.2 software was used for analysis of chromatograms.

## SUPPLEMENTARY MATERIALS

**Figure S1.**
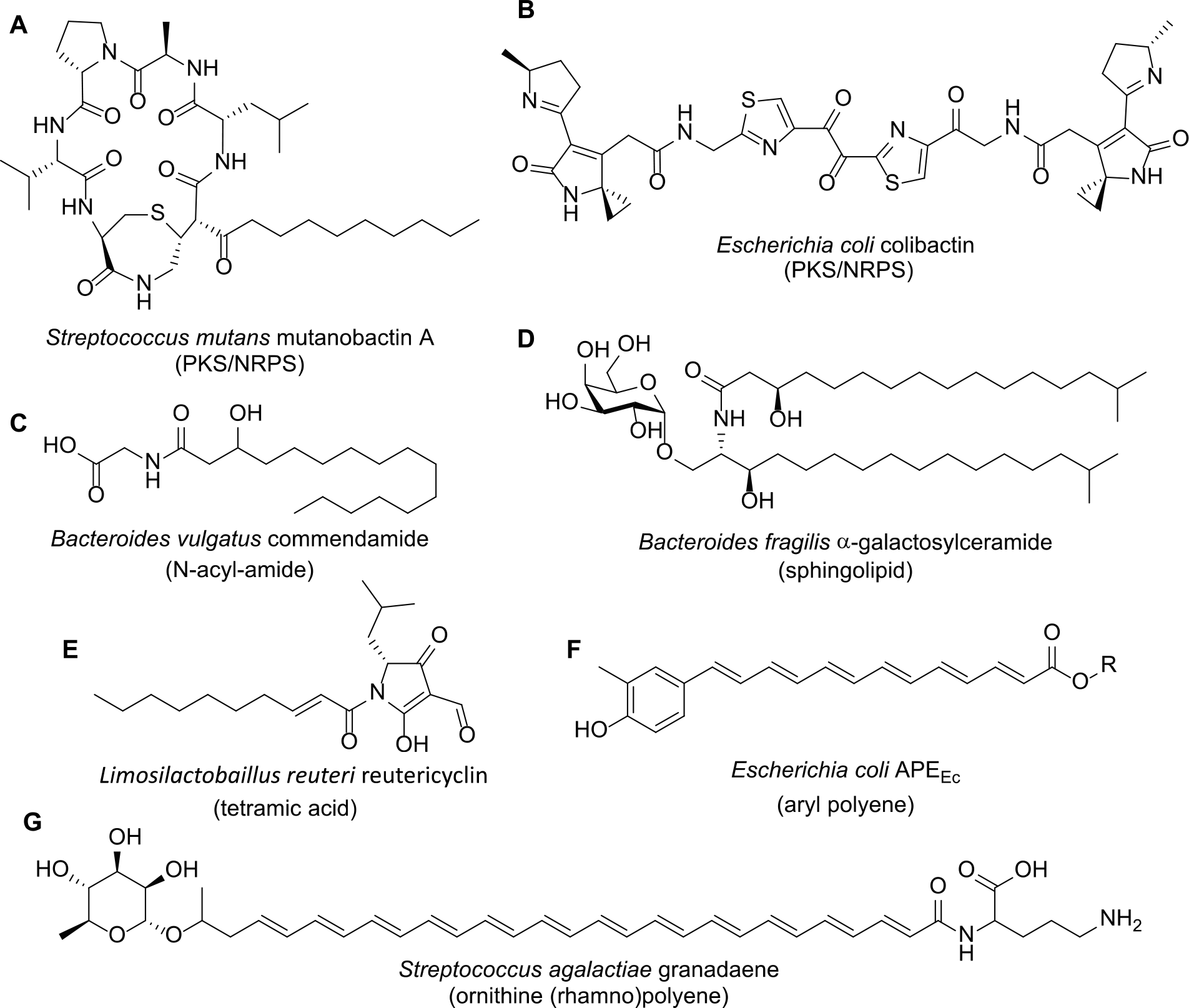
Structures of bioactive fatty acid-like, polyketide and NRPS/PKS hybrid compounds in bacteria from the gastrointestinal tract. So far, there are only seven microbial metabolites from GI tract have been chemically identified. These are an antifungal PKS/NRPS hybrid metabolite, Mutanobactin A produced by *Steptococcus mutans,* a genotoxin PKS/NRPS hybrid metabolite, Colibactin produced by *Escherichia coli*; a G-Protein Coupled Receptors (GPCR) agonist N-acyl-amide metabolite, Mutanobactin A, produced by *Bacteroides vulgatus*; an invariant Natural Killer Cell (iNTK) activator sphingolipid metabolite, α-galactosylceramide, produced by *Bacteroides fragilis*; an antimicrobial tetramic acid metabolite, Reutericyclin, produced by *Limosilactobacillus reuteri*; an antioxidant aryl polyene molecule, APEEC, produced by *Escherichia coli*; a polyene ornithine compound, Granadaene, produced by *Streptococcus agalactiae*. Related to Introduction.

**Figure S2.**
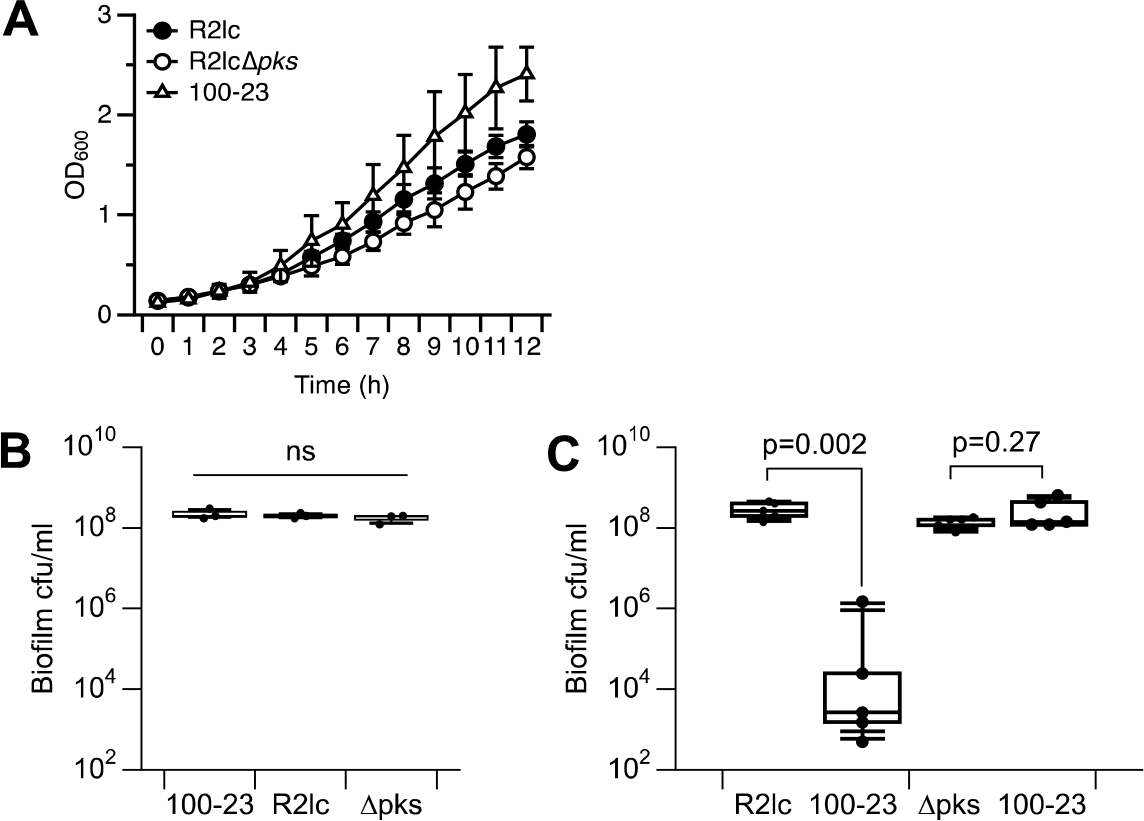
A polyketide synthase cluster in *L. reuteri* R2lc inhibits biofim formation of *L. reuteri* 100-23. A) R2lc and R2lcΔ*pks* have a similar growth dynamic, and 100-23 grows slightly faster compared to R2lc and R2lcΔ*pks* in mono-culture. B) Quantification of the number of CFU in single and C) co-culture biofilm. For box and whisker plots, the whiskers represent the maximum and minimal values, and the lower, middle and upper line of the box represent first quartile, median and third quartile, respectively. Circles represent individual data points. All data represent an average of at least three independent experiments. Statistical significance was determined by Wilcoxon / Kruskal-Wallis Tests p<0.05 considered as significant. p=P value. Related to Figure 1.

**Figure S3.**
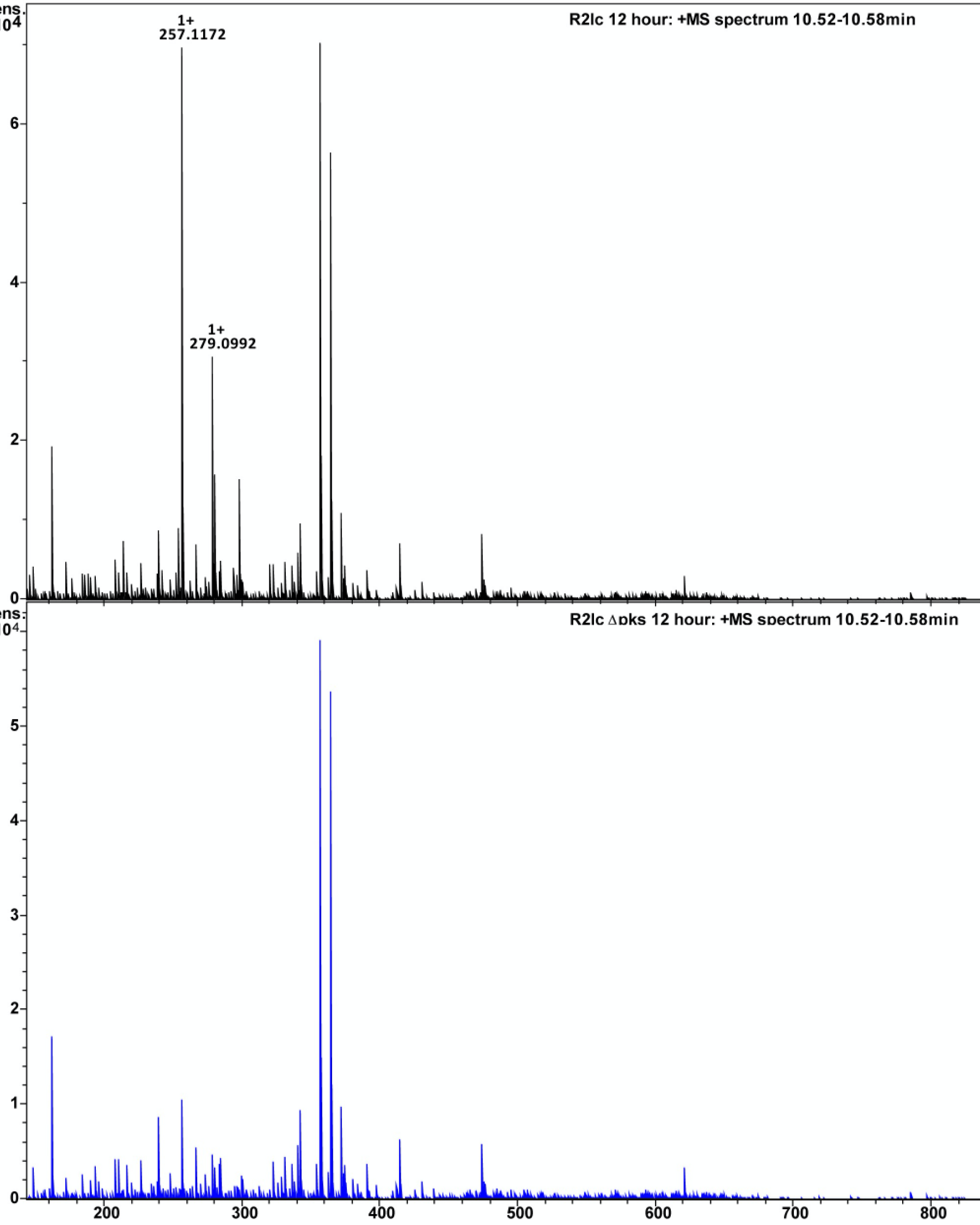
R2lc but not *pks* mutant produces unique compounds. The mass spectrum of the peak between 10.52 min –10.58 min in the R2lc chromatogram at 12 hours clearly showed an ion with m/zvalue of 257.1172 as the [M+H]+and 279.0092 as the corresponding [M+Na]+ which were absent in the corresponding 12-hour culture of Δpks. Bruker Smart Formula algorithm, which uses both the exact mass of the molecular ion and the isotopic pattern allowed accurate determination of the molecular formula as C16H16O3. Related to Figure 1.

**Figure S4.**
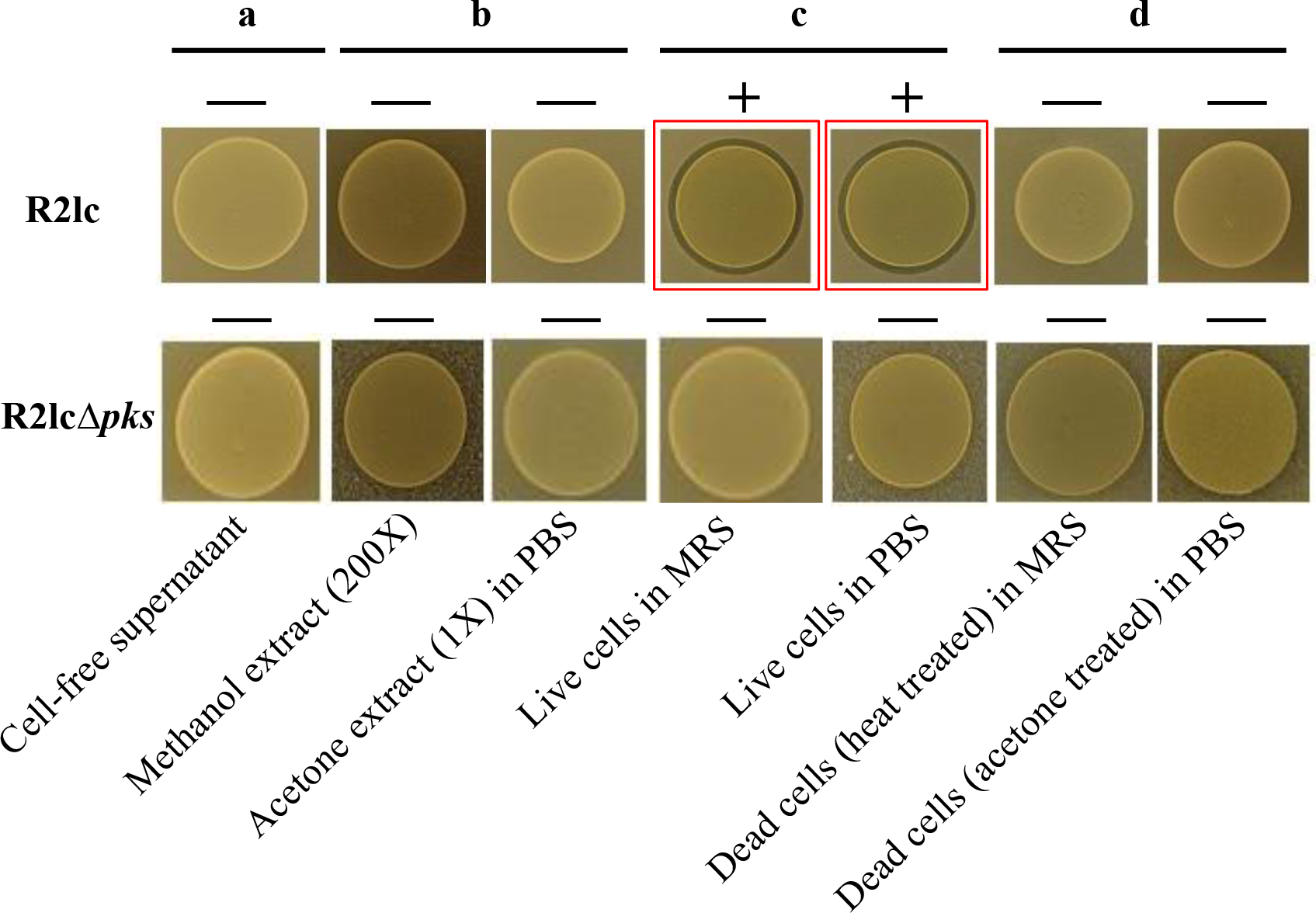
*Pks*-mediated killing mechanism requires Pks-producing strain to be alive. Double-layer agar diffusion assay with R2lc (top panel) or R2lcΔ*pks* (bottom panel) a) cell-free supernatants; b) concentrated methanol (which concentrates the pigment (Özçam et al., 2019) and acetone extracts; c) live cells in MRS and PBS; d) cells killed by treatment of heat or acetone. Related to Figure 1.

**Figure S5.**
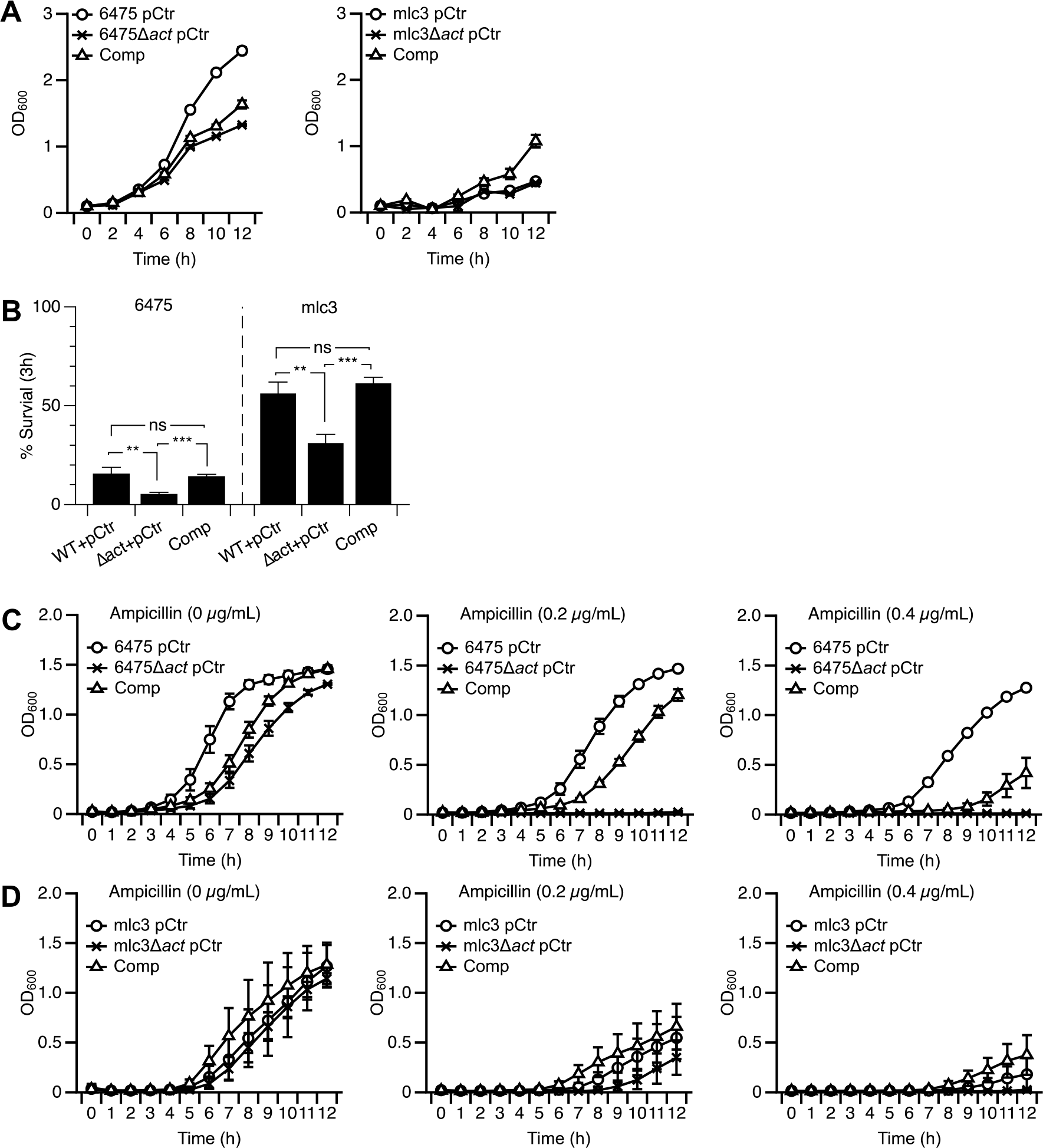
Growth dynamics of wild-type, *act* mutant and complemented strains of *L. reuteri* 6475 and mlc3 in MRS and MRS supplanted with lysozyme and ampicillin. A) Deletion and complementation of the *act* gene reduces the growth of *L. reuteri* 6475, while complementation but not deletion of the *act* gene increases the growth of *L. reuteri* mlc3. B) Deletion of the *act* gene in *L. reuteri* 6475 and mlc3 increases the lysozyme sensitivity, while the complemented strains survived at levels comparable to that of the wild-type strains. C) the *act* mutants of 6475 and D) mlc3 are more sensitive to lysozyme, and complemented strains had a phenotype similar to that of wild-type strains. Related to Figure 6.

**Table S1.** Strains used for comparative genome analyses.

| Strain Number | Strain | Sensitive/Resistant* |
| --- | --- | --- |
| 1 | CR | S |
| 2 | 100-23 | S |
| 3 | L1604-1 | S |
| 4 | ATCC53608 | S |
| 5 | 3c6 | S |
| 6 | L1600-1 | S |
| 7 | one-one | S |
| 8 | rat19 | S |
| 9 | JCM1081 | S |
| 10 | I5007 | S |
| 11 | 6799jm-1 | S |
| 12 | lpuph | S |
| 13 | CF48-3A1 | S |
| 14 | 1366 | S |
| 15 | N2J | S |
| 16 | LR4020 | S |
| 17 | SD2112 | S |
| 18 | AD23 | S |
| 19 | N4I | S |
| 20 | 100-93 | S |
| 21 | PTA6475 | R |
| 22 | DSM20016<sup>T</sup> | R |
| 23 | *mlc3* | R |
| 24 | Lr4000 | R |

\*Sensitive/resistant phenotypes were determined based on statistical analyses shown in Figure 4. S: sensitive, R: Resistant. Related to Figure 4.

**Table S2.** Unique genes in R2lc-resistat 6475, DSM20016^T^ and mlc3.

| Locus Tag | Gene Name | Length (aa) |
| --- | --- | --- |
| Lreu_0435 | hypothetical protein | 69 |
| Lreu_0848 | holin, Cph1 family | 149 |
| Lreu_1143 | transcription regulator, XRE family | 67 |
| Lreu_1353 | hypothetical protein | 522 |
| Lreu_1362 | glycosyltransferase | 324 |
| Lreu_1368 | acyltransferase | 355 |
| Lreu_1869 | hypothetical protein | 175 |

aa: amino acid. Related to Figure 4.

**Table S3.** Bacterial strains used in this study.

aa: amino acid. Related to Figure 4.
| Genus and Species | Strain | VPL identifier | Description/ Origin | Source |
| --- | --- | --- | --- | --- |
| *Escherichia coli* | EC1000 | VPL1009 | Cloning host | (Leenhouts et al., 1996) |
| *Lactococcus lactis* ssp. *cremoris* | MG1363 | VPL2005 | *L. lactis* MG1363 harboring pSIP411 | Laboratory stock |
| *Limosilactobaillus reuteri* | R2lc | VPL1053 | Rat | Siv Ahrné—JGI 2716884882 |
| *Limosilactobaillus reuteri* | R2lc::Cm | VPL4231 | Insertion of *Cm* gene in R2lc chromosome | This study |
| *Limosilactobaillus reuteri* | R2lcΔ*fun*::Cm | VPL4234 | Insertion of *Cm* gene in R2lcΔ*fun* chromosome | This study |
| *Limosilactobaillus reuteri* | R2lcΔ*pks*::Cm | VPL4209 | Insertion of *Cm* gene in R2lcΔ*pks* chromosome | (Özçam et al., 2019) |
| *Limosilactobaillus reuteri* | 6475 (pJP042) | VPL2047 | 6475 harboring pJP042 plasmid | (Van Pijkeren and Britton, 2012) |
| *Limosilactobaillus reuteri* | 6475 + pNZ8048 | VPL31613 | 6475—Rif<sup>R</sup> + pNZ8048 | This study |
| *Limosilactobaillus reuteri* | 6475Δ*act* | VPL4241 | Deletion of *act* gene in 6475 | This study |
| *Limosilactobaillus reuteri* | 6475Δ*act* + pNZ8048 | VPL31673 | 6475Δ*act*—Rif<sup>R</sup> + pNZ8048 | This study |
| *Limosilactobaillus reuteri* | 6475Δ*act*—Rif<sup>R</sup> (pSIP411) | VPL31153 | 6475Δ*act*—Rif<sup>R</sup> harboring pSIP411 plasmid | This study |
| *Limosilactobaillus reuteri* | 6475Δ*act*::*act*—Rif<sup>R</sup> | VPL4250 | 6475Δ*act*—Rif<sup>R</sup> harboring pSIP-*act* plasmid | This study |
| *Limosilactobaillus reuteri* | 6475Δ*act*::*act*—Rif<sup>R</sup> | VPL31664 | 6475 Δ*act* + pJP31601 | This study |
| *Limosilactobaillus reuteri* | 6475/leptin | VPL31313 | 6475 producing leptin | (Alexander et al., 2019) |
| *Limosilactobaillus reuteri* | 100-23 | VPL1049 | Rat | JGI: 2500069000 |
| *Limosilactobaillus reuteri* | 100-23—Rif<sup>R</sup> | VPL4251 | 100-23 Rif resistant | This study |
| *Limosilactobaillus reuteri* | 100-23/leptin | VPL315998 | 100-23—Rif<sup>R</sup> + pVPL31313 | This study |
| *Limosilactobaillus reuteri* | 100-23—Rif<sup>R</sup> (pSIP411) | VPL31155 | 100-23—Rif<sup>R</sup> harboring pSIP411 plasmid | This study |
| *Limosilactobaillus reuteri* | 100-23::act—Rif<sup>R</sup> | VPL4249 | Deletion of *act* gene in 100-23, Rif resistant | This study |
| *Limosilactobaillus reuteri* | 6475 | VPL1014 | Human | BioGaia AB |
| *Limosilactobaillus reuteri* | 6475—Rif<sup>R</sup> | VPL4296 | 6475, Rif resistant | This study |
| *Limosilactobaillus reuteri* | SD2112 | VPL1013 | Human | BioGaia AB |
| *Limosilactobaillus reuteri* | SD2112—Rif<sup>R</sup> | VPL4297 | SD2112, Rif resistant | This study |
| *Limosilactobaillus reuteri* | CSB7 | VPL1168 | Chicken | Jens Walter |
| *Limosilactobaillus reuteri* | CSB7—Rif<sup>R</sup> | VPL4298 | CSB7, Rif resistant | This study |
| *Limosilactobaillus reuteri* | KYE26 | VPL1162 | Chicken | Jens Walter |
| *Limosilactobaillus reuteri* | KYE26—Rif<sup>R</sup> | VPL4299 | KYE26, Rif resistant | This study |
| *Limosilactobaillus reuteri* | LK159 | VPL1156 | Chicken | Jens Walter |
| *Limosilactobaillus reuteri* | LK159—Rif<sup>R</sup> | VPL4301 | LK159, Rif resistant | This study |
| *Limosilactobaillus reuteri* | L4 | VPL1173 | Chicken | Jens Walter |
| *Limosilactobaillus reuteri* | L4—Rif<sup>R</sup> | VPL4302 | L4, Rif resistant | This study |
| *Limosilactobaillus reuteri* | mouse2 | VPL1070 | Mouse | (Frese et al., 2011) |
| *Limosilactobaillus reuteri* | Mouse2—Rif<sup>R</sup> | VPL4252 | Mouse2, Rif resistant | This study |
| *Limosilactobaillus reuteri* | N2J | VPL1052 | Rat | (Frese et al. 2011) |
| *Limosilactobaillus reuteri* | N2J—Rif<sup>R</sup> | VPL4253 | N2J, Rif resistant | This study |
| *Limosilactobaillus reuteri* | one-one | VPL1060 | Mouse | (Frese et al. 2011) |
| *Limosilactobaillus reuteri* | one-one—Rif<sup>R</sup> | VPL4254 | one-one, Rif resistant | This study |
| *Limosilactobaillus reuteri* | ATCC 53608 | VPL1090 | Pig | BioGaia AB EMBL LN906634 |
| *Limosilactobaillus reuteri* | ATCC 53608—Rif<sup>R</sup> | VPL4255 | ATCC 53608, Rif resistant | This study |
| *Limosilactobaillus reuteri* | L1600-1 | VPL1064 | Mouse | (Frese et al. 2011) |
| *Limosilactobaillus reuteri* | L1600-1—Rif<sup>R</sup> | VPL4256 | L1600-1, Rif resistant | This study |
| *Limosilactobaillus reuteri* | N2D | VPL1067 | Rat | Siv Ahrné |
| *Limosilactobaillus reuteri* | N2D—Rif<sup>R</sup> | VPL4257 | N2D, Rif resistant | This study |
| *Limosilactobaillus reuteri* | DSM17938 | VPL4171 | Human | BioGaia AB |
| *Limosilactobaillus reuteri* | DSM17938—Rif<sup>R</sup> | VPL4258 | DSM17938, Rif resistant | This study |
| *Limosilactobaillus reuteri* | DSM17938Δact | VPL4313 | Deletion of *act* gene in DSM17938 | This study |
| *Limosilactobaillus reuteri* | DSM17938Δact—Rif<sup>R</sup> | VPL4315 | DSM17938Δact, Rif resistant | This study |
| *Limosilactobaillus reuteri* | Lr4000 | VPL1071 | Mouse | BioGaia AB |
| *Limosilactobaillus reuteri* | Lr4000—Rif<sup>R</sup> | VPL4259 | Lr4000, Rif resistant | This study |
| *Limosilactobaillus reuteri* | CF48-3A1 | VPL1086 | Human | BioGaia AB—JGI2502171173 |
| *Limosilactobaillus reuteri* | CF48-3A1—Rif<sup>R</sup> | VPL4260 | CF48-3A1, Rif resistant | This study |
| *Limosilactobaillus reuteri* | L1604-1 | VPL1066 | Mouse | (Frese et al. 2011) |
| *Limosilactobaillus reuteri* | L1604-1—Rif<sup>R</sup> | VPL4261 | L1604-1, Rif resistant | This study |
| *Limosilactobaillus reuteri* | CR | VPL1059 | Rat | (Frese et al. 2011) |
| *Limosilactobaillus reuteri* | CR—Rif<sup>R</sup> | VPL4262 | CR, Rif resistant | This study |
| *Limosilactobaillus reuteri* | mlc3 | VPL1050 | Mouse | JGI: 2506381016 |
| *Limosilactobaillus reuteri* | mlc3—Rif<sup>R</sup> | VPL4263 | mlc3, Rif resistant | This study |
| *Limosilactobaillus reuteri* | mlc3 + pNZ8048 | Vpl31668 | VPL4263 + pNZ8048 | This study |
| *Limosilactobaillus reuteri* | mlc3Δact | VPL4246 | Deletion of *act* gene in mlc3 | This study |
| *Limosilactobaillus reuteri* | mlc3Δact—Rif<sup>R</sup> | VPL4308 | mlc3Δact, Rif resistant | This study |
| *Limosilactobaillus reuteri* | mlc3Δact + pNZ8048 | VPL31675 | VPL4308+ pNZ8048 | This study |
| *Limosilactobaillus reuteri* | mlc3Δact—Rif<sup>R</sup> (pSIP411) | VPL31177 | mlc3Δact—Rif<sup>R</sup>, harboring pSIP411 plasmid | This study |
| *Limosilactobaillus reuteri* | mlc3Δact::act—Rif<sup>R</sup> | VPL4248 | mlc3Δact—Rif<sup>R</sup> harboring pSIP-act plasmid | This study |
| *Limosilactobaillus reuteri* | mlc3Δact::act—Rif<sup>R</sup> | VPL31666 | mlc3Δact + pVPL31603 | This study |
| *Limosilactobaillus reuteri* | I5007 | VPL1082 | Pig | JGI: 2554235423 |
| *Limosilactobaillus reuteri* | I5007—Rif<sup>R</sup> | VPL4264 | I5007, Rif resistant | This study |
| *Limosilactobaillus reuteri* | Lr4020 | VPL1072 | Mouse | (Frese et al. 2011) |
| *Limosilactobaillus reuteri* | Lr4020—Rif<sup>R</sup> | VPL4265 | Lr4020, Rif resistant | This study |
| *Limosilactobaillus reuteri* | Lp167-67 | VPL1085 | Pig | BioGaia AB JGI 2599185361 |
| *Limosilactobaillus reuteri* | Lp167-67—Rif<sup>R</sup> | VPL4266 | Lp167-67, Rif resistant | This study |
| *Limosilactobaillus reuteri* | N4I | VPL1063 | Rat | (Frese et al. 2011) |
| *Limosilactobaillus reuteri* | N4I—Rif<sup>R</sup> | VPL4267 | N4I, Rif resistant | This study |
| *Limosilactobaillus reuteri* | 3c6 | VPL1083 | Pig | JGI: 2599185333 |
| *Limosilactobaillus reuteri* | 3c6—Rif<sup>R</sup> | VPL4268 | 3c6, Rif resistant | This study |
| *Limosilactobaillus reuteri* | FUA3043 | VPL1062 | Rat | (Frese *et al.* 2011) |
| *Limosilactobaillus reuteri* | FUA3043—Rif<sup>R</sup> | VPL4269 | FUA3043, Rif resistant | This study |
| *Limosilactobaillus reuteri* | Rat 19 | VPL1069 | Rat | (Frese *et al.* 2011) |
| *Limosilactobaillus reuteri* | Rat 19—Rif<sup>R</sup> | VPL4270 | Rat 19, Rif resistant | This study |
| *Limosilactobaillus reuteri* | 6799jm-1 | VPL1051 | Mouse | (Frese *et al.* 2011) |
| *Limosilactobaillus reuteri* | 6799jm-1—Rif<sup>R</sup> | VPL4271 | 6799jm-1, Rif resistant | This study |
| *Limosilactobaillus reuteri* | 100-93 | VPL1047 | Mouse | (Frese *et al.* 2011) |
| *Limosilactobaillus reuteri* | 100-93—Rif<sup>R</sup> | VPL4272 | 100-93, Rif resistant | This study |
| *Limosilactobaillus reuteri* | Lpuph-1 | VPL1056 | Mouse | JGI: 2506381017 |
| *Limosilactobaillus reuteri* | Lpuph-1—Rif<sup>R</sup> | VPL4273 | Lpuph-1, Rif resistant | This study |
| *Limosilactobaillus reuteri* | DSM 20016 | VPL1046 | Human | JGI: 640427118 |
| *Limosilactobaillus reuteri* | DSM 20016—Rif<sup>R</sup> | VPL4274 | DSM 20016, Rif resistant | This study |
| *Limosilactobaillus reuteri* | AD 23 | VPL1048 | Rat | (Frese *et al.* 2011) |
| *Limosilactobaillus reuteri* | AD 23—Rif<sup>R</sup> | VPL4275 | AD 23, Rif resistant | This study |
| *Limosilactobaillus reuteri* | 173/4 | VPL1135 | Pig | Jens Walter |
| *Limosilactobaillus reuteri* | 173/4—Rif<sup>R</sup> | VPL4276 | 173/4, Rif resistant | This study |
| *Limosilactobaillus reuteri* | 4S17 | VPL1122 | Pig | (Frese *et al.* 2011) |
| *Limosilactobaillus reuteri* | 4S17—Rif<sup>R</sup> | VPL4277 | 4S17, Rif resistant | This study |
| *Limosilactobaillus reuteri* | 1366 | VPL1115 | Chicken | Jens Walter |
| *Limosilactobaillus reuteri* | 1366—Rif<sup>R</sup> | VPL4278 | 1366, Rif resistant | This study |
| *Limosilactobaillus reuteri* | JW2015 | VPL1126 | Pig | Jens Walter |
| *Limosilactobaillus reuteri* | JW2015—Rif<sup>R</sup> | VPL4279 | JW2015, Rif resistant | This study |
| *Limosilactobaillus reuteri* | 32 | VPL1137 | Pig | Jens Walter |
| *Limosilactobaillus reuteri* | 32—Rif<sup>R</sup> | VPL4280 | 32, Rif resistant | This study |
| *Limosilactobaillus reuteri* | 3S3 | VPL1120 | Pig | Jens Walter |
| *Limosilactobaillus reuteri* | 3S3—Rif<sup>R</sup> | VPL4281 | 3S3, Rif resistant | This study |
| *Limosilactobaillus reuteri* | 13S14 | VPL1118 | Pig | Jens Walter |
| *Limosilactobaillus reuteri* | 13S14—Rif<sup>R</sup> | VPL4282 | 13S14, Rif resistant | This study |
| *Limosilactobaillus reuteri* | LK150 | VPL1112 | Chicken | Jens Walter |
| *Limosilactobaillus reuteri* | LK150—Rif<sup>R</sup> | VPL4283 | LK150, Rif resistant | This study |
| *Limosilactobaillus reuteri* | JCM 1081 | VPL1110 | Chicken | Jens Walter |
| *Limosilactobaillus reuteri* | JCM 1081—Rif<sup>R</sup> | VPL4284 | JCM 1081, Rif resistant | This study |
| *Limosilactobaillus reuteri* | HW8 | VPL1166 | Chicken | Jens Walter |
| *Limosilactobaillus reuteri* | HW8—Rif<sup>R</sup> | VPL4300 | HW8, Rif resistant | This study |
| *Limosilactobaillus reuteri* | JW2019 | VPL1129 | Pig | Jens Walter |
| *Limosilactobaillus reuteri* | JW2019—Rif<sup>R</sup> | VPL4286 | JW2019, Rif resistant | This study |
| *Limosilactobaillus reuteri* | 1133 (146/2) | VPL1133 | Pig | Jens Walter |
| *Limosilactobaillus reuteri* | 1133 (146/2)—Rif<sup>R</sup> | VPL4287 | 1133, Rif resistant | This study |
| *Limosilactobaillus reuteri* | CP415 | VPL1146 | Pig | Jens Walter |
| *Limosilactobaillus reuteri* | CP415—Rif<sup>R</sup> | VPL4288 | CP415, Rif resistant | This study |
| *Limosilactobaillus reuteri* | 1063 | VPL1143 | Pig | Jens Walter |
| *Limosilactobaillus reuteri* | 1063—Rif<sup>R</sup> | VPL4289 | 1063, Rif resistant | This study |
| *Limosilactobaillus reuteri* | 1068 | VPL1142 | Pig | Jens Walter |
| *Limosilactobaillus reuteri* | 1068—Rif<sup>R</sup> | VPL4291 | 1068, Rif resistant | This study |
| *Limosilactobaillus reuteri* | 1048 | VPL1141 | Pig | Jens Walter |
| *Limosilactobaillus reuteri* | 1048—Rif<sup>R</sup> | VPL4292 | 1048, Rif resistant | This study |
| *Limosilactobaillus reuteri* | 1013 | VPL1140 | Pig | Jens Walter |
| *Limosilactobaillus reuteri* | 1013—Rif<sup>R</sup> | VPL4293 | 1013, Rif resistant | This study |
| *Limosilactobaillus reuteri* | 1704 | VPL1139 | Pig | Jens Walter |
| *Limosilactobaillus reuteri* | 1704—Rif<sup>R</sup> | VPL4294 | 1704, Rif resistant | This study |
| *Limosilactobaillus reuteri* | FUA3048 | VPL1063 | Rat | (Frese *et al.* 2011) |
| *Limosilactobaillus reuteri* | FUA3048 —Rif<sup>R</sup> | VPL4295 | FUA3048, Rif resistant | This study |

VPL: Van Pijkeren Laboratory strain identification number. pVPL: Van Pijkeren Laboratory plasmid identification number. Rif<sup>R</sup>: rifampicin-resistant; *act*: acyltransferase; JGI: Joint Genome Institute; Cm: Chloramphenicol; Rif: Rifampicin.

**Table S4.**
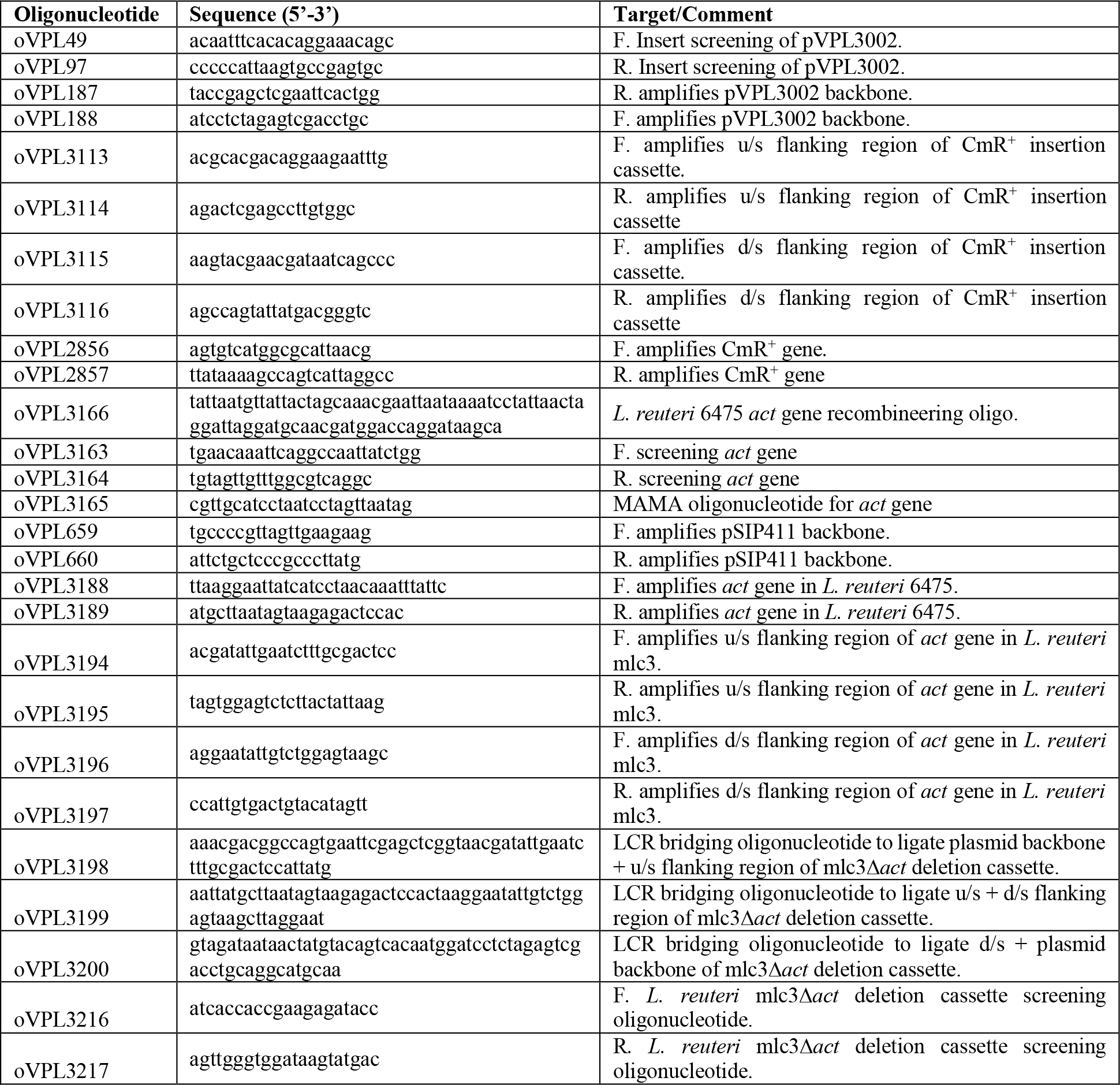
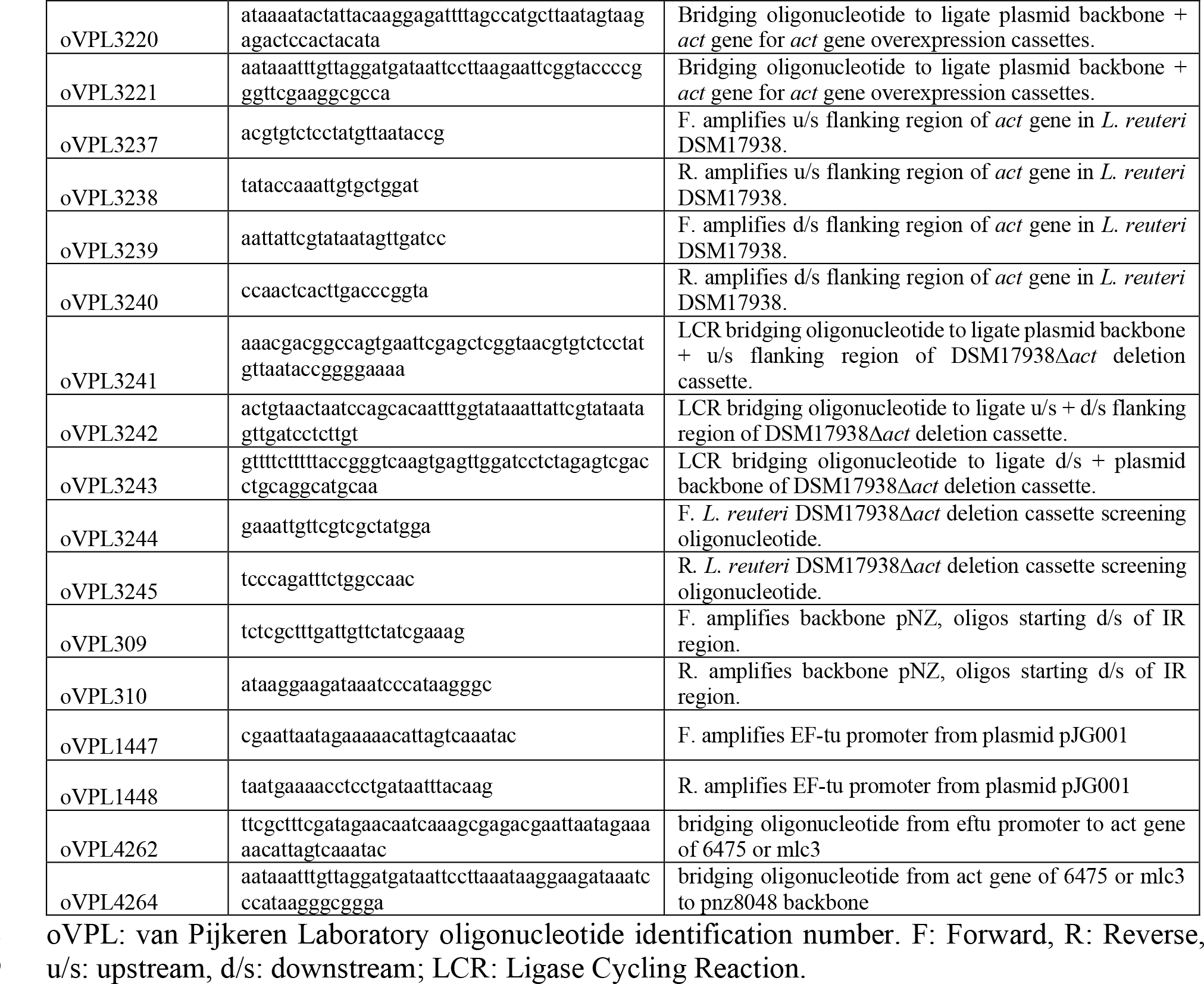
Oligonucleotides used in this study.

**Table S5.** Plasmids used in this study.

| Plasmid | Characteristic | Source |
| --- | --- | --- |
| pSIP411 | EmR, Sakacin-P based expression vector | (Sørvig et al., 2005) |
| pVPL3002 | pORI19 harboring *L. reuteri* derived ddlF258Y | (Zhang et al., 2018) |
| pVPL31150 | pSIP411 harboring *L. reuteri* 6475 *act* gene | This Study |
| pVPL2042 | Em<sup>R</sup>, pNZ8048 derivative. Cm marker was replaced by Em marker | (Zhang et al., 2018) |
| pVPL31130 | Em<sup>R</sup>, derivative of vector pVPL3002 in which the *L. reuteri*<br>R2lc::Cm and R2lcΔpks::Cm insertion cassette was cloned in the<br>MCS site. | This Study |
| pVPL31079 | Em<sup>R</sup>, derivative of vector pVPL3002 in which the *L. reuteri*<br>mlc3Δ*act* deletion cassette was cloned in the MCS site. | This Study |
| pVPL31139 | Em<sup>R</sup>, derivative of vector pVPL3002 in which the *L. reuteri*<br>DSM17938Δ*act* deletion cassette was cloned in the MCS site. | This Study |
| pVPL31601 | Em<sup>R</sup>, derivative of vector pVPL3002 in which the *L. reuteri* mlc3<br>*act* gene was cloned in the MCS site. | This Study |
| pVPL31603 | Em<sup>R</sup>, derivative of vector pVPL3002 in which the *L. reuteri*<br>ATCC6475 *act* gene was cloned in the MCS site. | This Study |

pVPL: Van Pijkeren Laboratory plasmid identification number. Em<sup>R</sup>: Erythromycin-resistant; Cm<sup>R</sup>: Chloramphenicol-resistant. Related to STAR Methods.

**Table S6.**
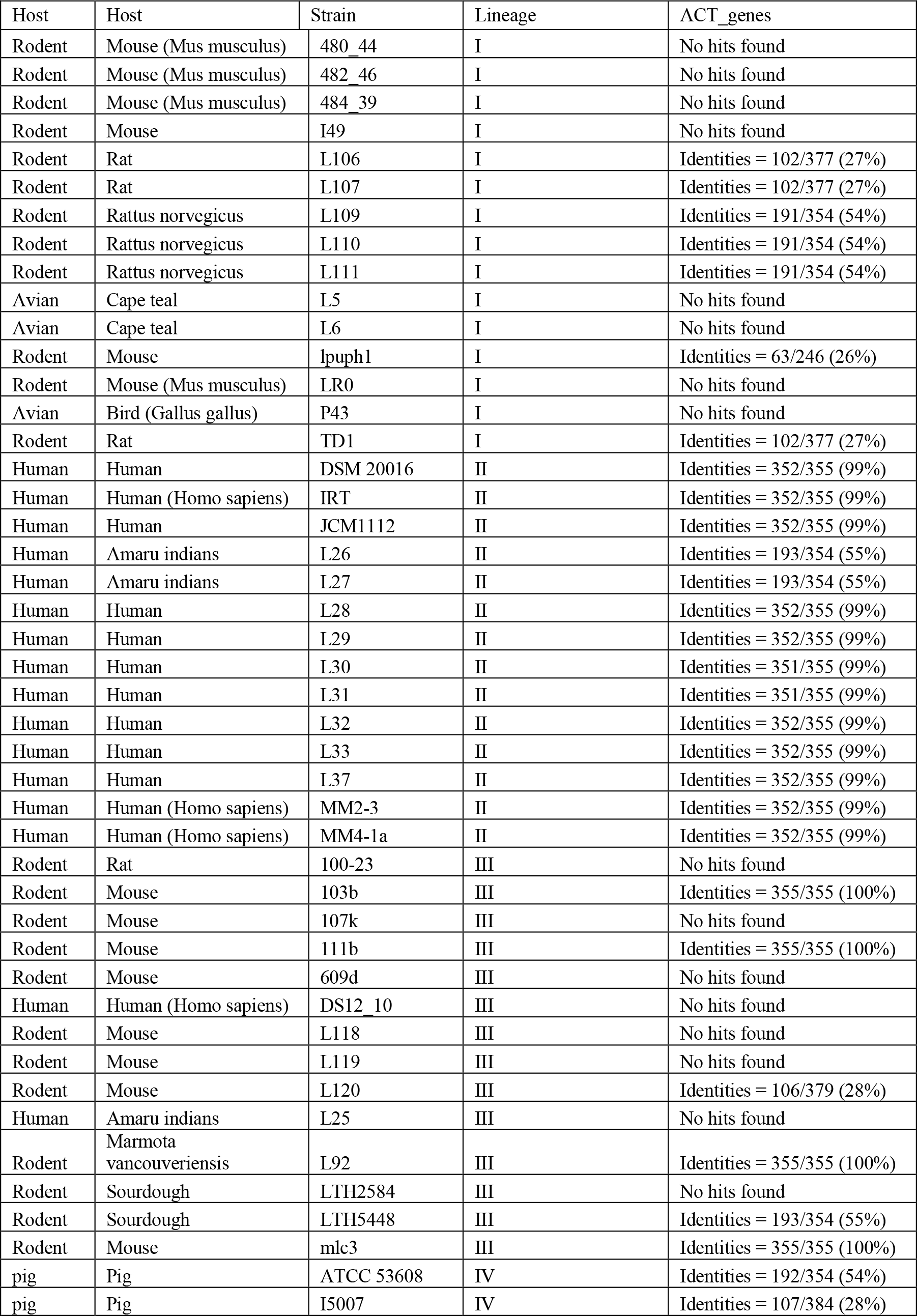
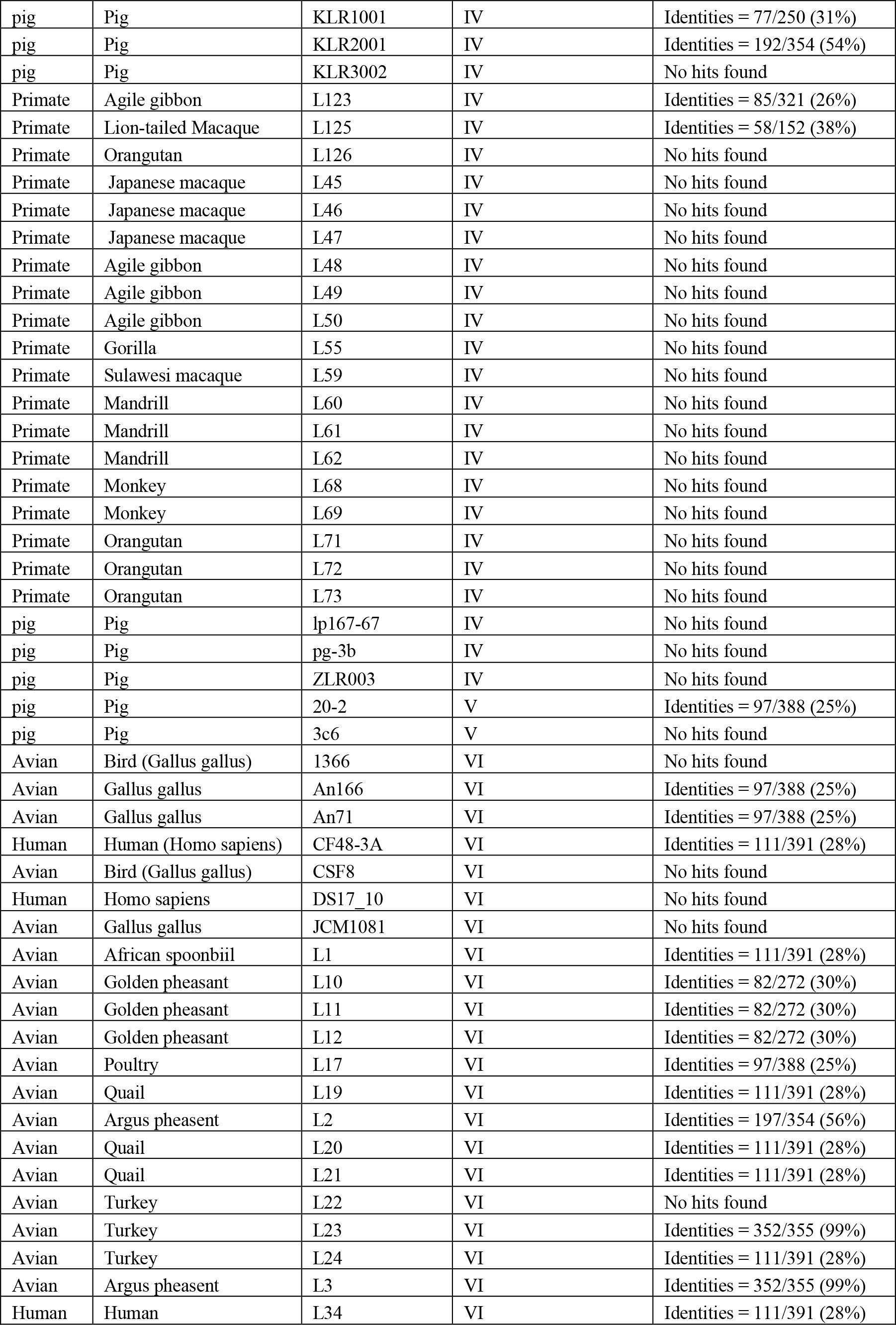
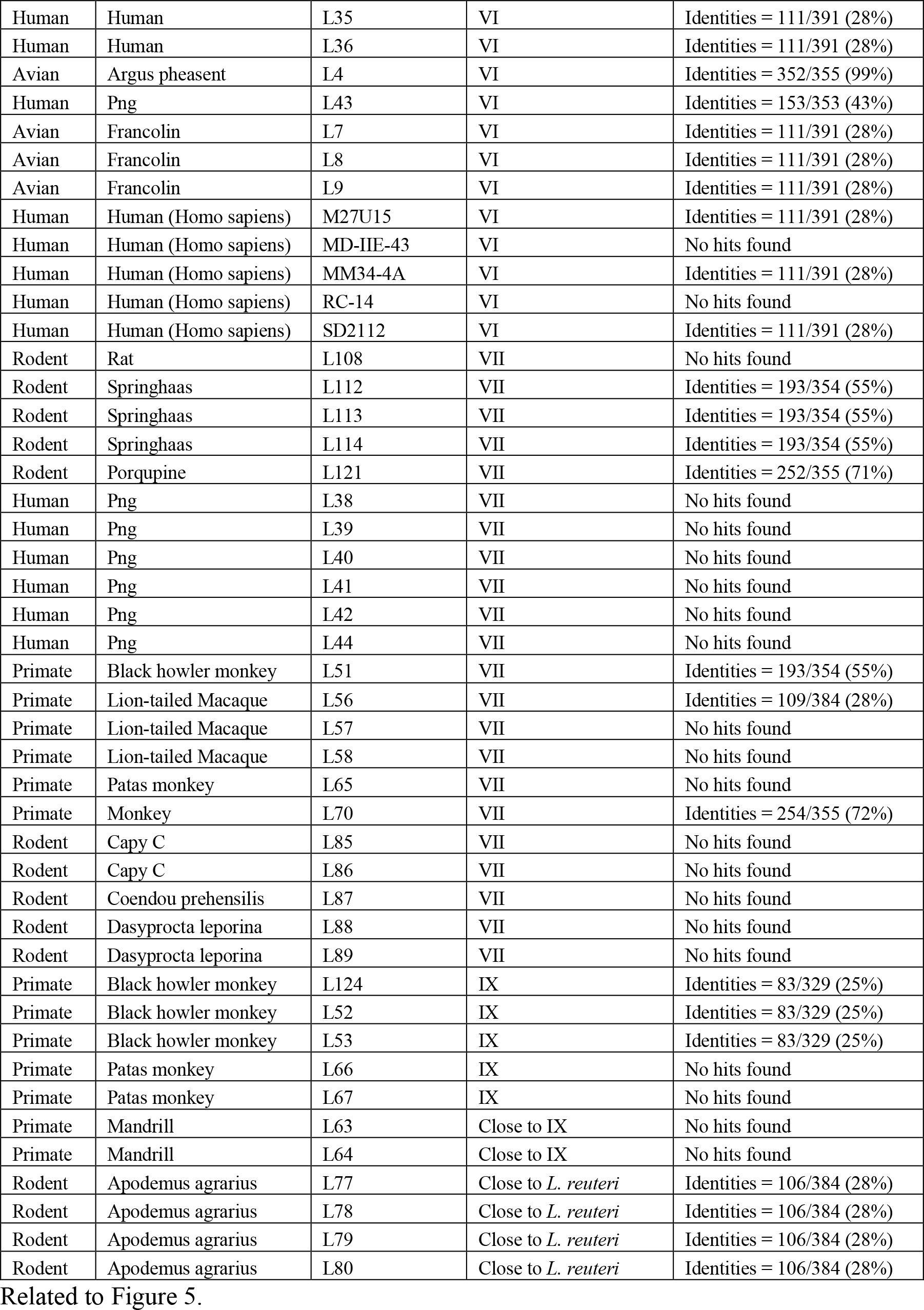
Acyltransferase gene in *L. reuteri* strains

